# Structural insights into wiring specificity in the neuromuscular system through the Beat-Side complex

**DOI:** 10.1101/2025.06.05.656487

**Authors:** Jessica M. Priest, Ruiling Zhang, Agnieszka M. Olechwier, Asaf B. Caspi-Lebovic, James Ashley, Vasudha Aher, Robert A. Carrillo, Engin Özkan

**Author notes:** These authors contributed equally (Jessica M. Priest, Ruiling Zhang, Agnieszka M. Olechwier, Asaf B. Caspi-Lebovic).

## Abstract

Nervous system assembly is guided by the actions of cell surface receptors. In *Drosophila*, members of the Beaten Path (Beat) and Sidestep (Side) protein families have been described as axon guidance receptor-cue pairs, in addition to roles in specifying synaptic connectivity in the optic lobe. To understand the molecular basis and specificity of Beat-Side interactions, we report here the first Beat-Side structure, Beat-Vc bound to Side-VI. The structure showed a binding topology similar to other neuronal immunoglobulin superfamily receptors, especially Nectins, SynCAMs, Dprs and DIPs, despite lack of established evolutionary relationships. Using a structure-based rational approach, we engineered and validated point mutations to break the binding between Beats and Sides. Using these mutant variants, we demonstrated in developing *Drosophila* larvae that the interaction between Beat-Ia and Side is required for establishing proper connectivity of motor neurons with muscles.

## INTRODUCTION

The development of the neuromuscular system is dependent on a molecular landscape controlled by cell-surface and secreted proteins. Motor axons rely on these proteins to mediate various interactions with the surrounding tissue,^1^ guide their movements, trigger cell adhesion, innervate muscles precisely, and form synaptic connections.^2,3^ Correct motor neuron-muscle pairing underlies all locomotor behaviors.

In the *Drosophila* embryo, motor neurons extend axonal projections from the ventral nerve cord to the periphery and innervate proper muscles through the combinatorial actions of guidance cue-receptor complexes and cell adhesion molecules.^1^ Two genes, Beaten Path (Beat) and Sidestep (Side), were identified in genetic screens three decades ago, and loss of these genes causes very similar and highly penetrant defects in motor axon guidance.^4–6^ Beat is expressed on growing motor axons, while Side is dynamically expressed on substrate tissue as the Beat-expressing motor axons elongate.^6,7^ This attractive guidance system allows for motor axons to branch and leave nerve bundles, counteracting adhesive forces mediated by cell adhesion molecules, such as the NCAM ortholog Fasciclin 2.^5,8^

Recent work has revealed that Beat and Side physically interact, likely functioning as a classical attractive guidance receptor-cue pair.^7^ Furthermore, Beat and Side were discovered to be members of two protein families, with 14 and 8 closely related members, respectively.^9–11^ High-throughput interaction screening uncovered a set of interactions between specific Beat-Side pairs,^12,13^ suggesting a model where neuronal Beat and Side expression patterns may direct the formation of circuits in a Beat-Side cognate pair-dependent manner. In support of this model, several Beats and Sides were observed to be expressed in a stereotyped manner in fly optic lobe neurons,^14,15^ and the Side-II–Beat-VI pair was shown to mediate wiring specificity between pre- and post-synaptic partner neurons, respectively, in the directionally selective motion detection circuit.^16^ Recent work demonstrated a role for the Side-IV–Beat-IIb cognate pair in synapse formation and localization in the visual system.^17^

Despite three decades of functional characterization and established roles in neuronal wiring, the structural details of Beats, Sides and their complexes remain elusive, and it is unknown how Beats and Sides achieve binding specificity with their binding partners. Here, we report the first structure of a Beat bound to a Side. Our structure revealed the interactions between the N-terminal domains of Beat-Vc and Side-VI, and allowed us to analyze the interface for determinants of specificity between Beat–Side pairs. We confirmed our structural insights by engineering mutant Beats and Sides that break their complexes. Finally, we showed that fruit flies with *beat-Ia* and *side* point mutants engineered to break their interaction show strong motor axon branching and targeting defects in fly larvae, validating our biochemical insights *in vivo*.

## RESULTS

### The first IG domains of Beat-Vc and Side-VI interact with each other

The 14 Beats and 8 Sides in *D. melanogaster* are cell surface receptors belonging to the immunoglobulin superfamily (IgSF). Beats invariably contain two extracellular immunoglobulin (IG) domains in their N-termini, followed by a transmembrane (TM) helix or GPI anchor. Sides have five IG domains followed by a Fibronectin-type III (FN) domain, a transmembrane helix, and a 200-400-residue intracellular region, predicted as unstructured, ending in a C-terminal PDZ peptide (Figure 1A). We first set out to show which domains are involved in Beat–Side interactions. For this, we employed the high-throughput extracellular interactome assay (ECIA), which was first used to reveal that multiple Beats and Sides form complexes with each other.^12^ ECIA depends on the expression of ectodomains in ‘bait’ and ‘prey’ formats in oligomerized form to mimic the clustered cell adhesion molecule complexes while increasing sensitivity via avidity (Figure 1B). We observed that the binding between the founding members of the families, Beat-Ia and Side, could be observed only when the first two IG domains of Beat-Ia and the first IG domain of Side were present, regardless of whether we use Beat-Ia or Side as bait or prey (Figure 1C and S1). Recent work has now shown definitively that the first IG domains of Beat-Ia and Side are necessary and sufficient to form complexes,^18^ which is in agreement with our results.

**Figure 1.**
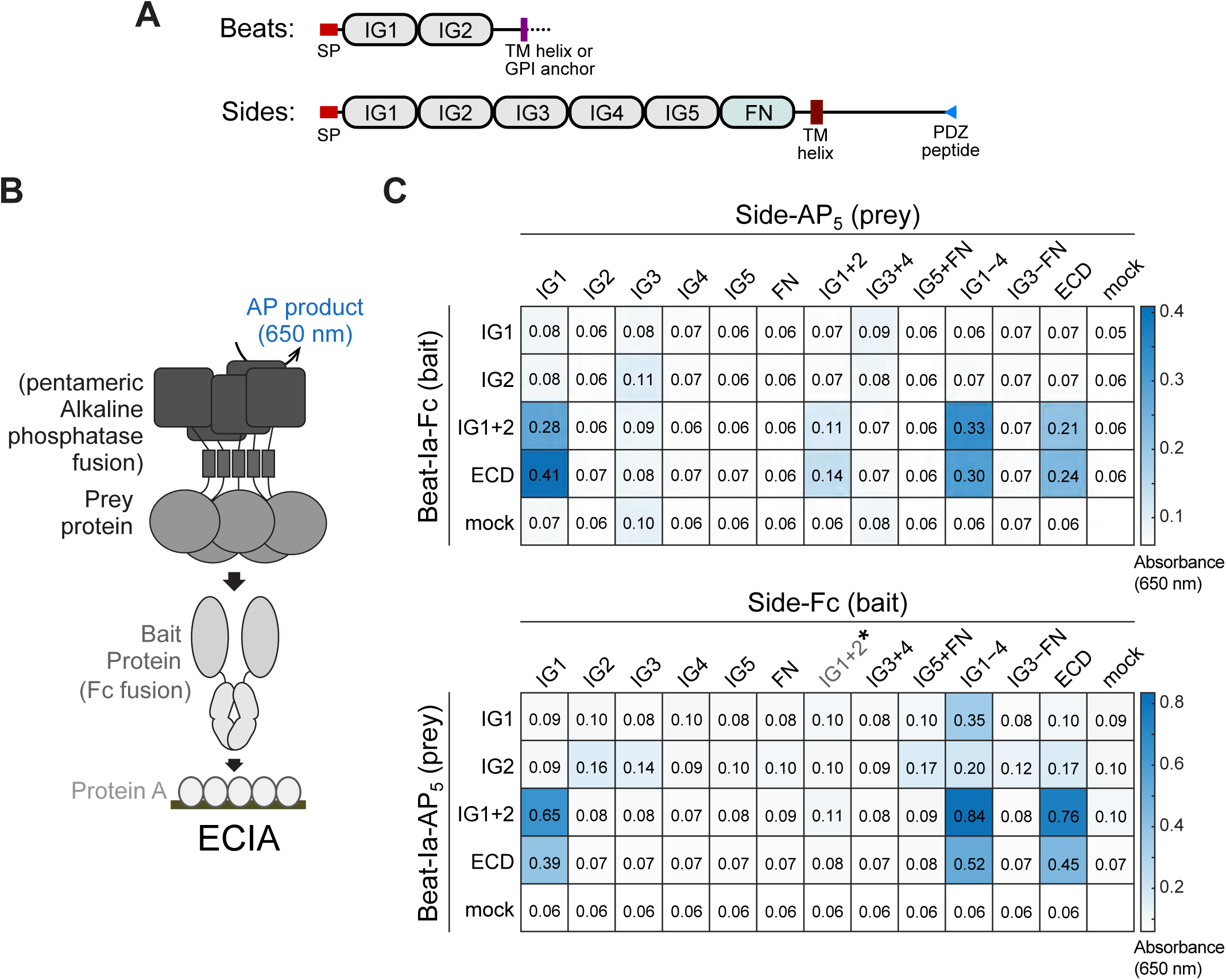
Pairwise interactions between Beat-Ia and Side domains. **A.** Domain organization of Beat and Side family proteins. SP: Signal Peptide. **B.** The design of ECIA. **C.** ECIA for Beat-Ia and Side domain-based constructs. The assay was duplicated with Beat-Ia as bait (top) and Side as bait (bottom). The asterisk indicates no expression for the construct. Western blots showing expression levels for all proteins are in Figure S1.

To gain insight into the molecular details of how Beats and Sides interact, we set out on a crystallization campaign of several Beats, Sides and their complexes. We determined the crystal structures of the two IG domains of Beat-Vb, the first IG domains of Side-IV, Side-VI and Side-VII, and the complex of Beat-Vc IG1+2 domains bound to Side-VI IG1 domain at 3.9 Å resolution (Table S1). The asymmetric unit of the complex crystals had two copies of Beat-Vc and Side-VI, each (Figures 2A), and we observed two plausible interaction interfaces based on the structural model (Figure 2B). The first contact interface buried a surface area of 1565 Å^2^, with 8 H-bonds and 3+ salt bridges present (Figures 2C-D), while the second interface was much smaller (buried surface area, 995 Å^2^) with only a few non-covalent contacts (Figure 2E).

**Figure 2.**
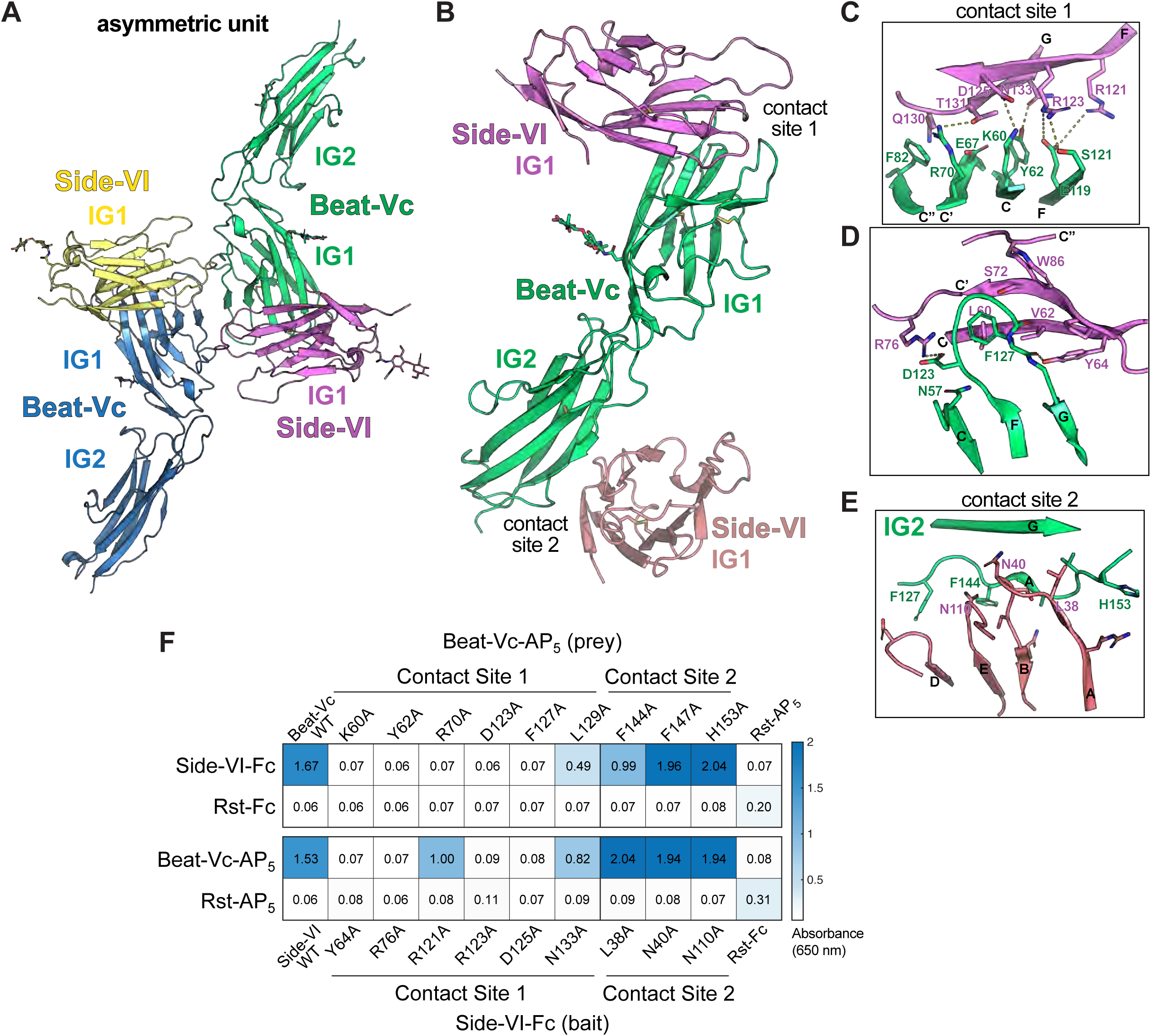
Crystal structure of the Beat-Vc–Side-VI complex. **A.** The asymmetric unit of the Beat-Vc IG1+2–Side-VI IG1 crystals contain two copies of Beat and Side each. N-linked glycans are shown as sticks. **B.** The two Beat–Side contacts observed in the crystals. The magenta and brown copies of Side-VI are crystallographic symmetry-related copies. **C,D.** Close view of contacts at the first contact site. Dashes indicate hydrogen bonds or salt bridges. **E.** Close view of contacts at the second contact site. **F.** ECIA with Beat-Vc and Side-VI ectodomains mutated at the first and second contact sites, which shows that Beat-Vc–Side-VI binding requires contacts only at the first contact site. See Figure S2 for western blots for the mutant proteins expressed.

We mutagenized both contact sites on both the Beat-Vc and the Side-VI surfaces and showed using ECIA that mutations at the first contact site either diminished or abolished Beat-Vc–Side-VI binding, while mutations at the second site had no significant effect on Beat-Vc–Side-VI complex formation (Figures 2F and S2), showing that the complex created by the first contact site may represent the physiological structure, while the second contact site is only a crystal contact. The validated complex shows interactions only between the first IG domains of Beat-Vc and Side-VI.

### Comparison of the Beat-Vc–Side-VI complex with known adhesive IG-IG complexes

The Beat-Vc–Side-VI complex structure shows that the *CCʹCʺFG* faces of the first domains of Beat-Vc and Side-VI interact directly. This is reminiscent of several neuronal IG-IG complexes, where the first domains mediate trans-cellular interactions (Figure 3A). Structure-based search of known structures with our Beat-Vc–Side-VI model returned the Nectin dimers as most similar, with a root-mean squared deviation (RMSD) of 4 Å to the Nectin1 homodimer (PDB:3ALP), and has weaker similarity to Dpr–DIP complexes as well (Figures 3A-B). This mode of IG1-IG1 interaction mediated by the *CCʹCʺFG* sheet appears to impose a common adhesion complex topology to IgSF adhesion molecules on neuronal surfaces, often at or near the synapse.

**Figure 3.**
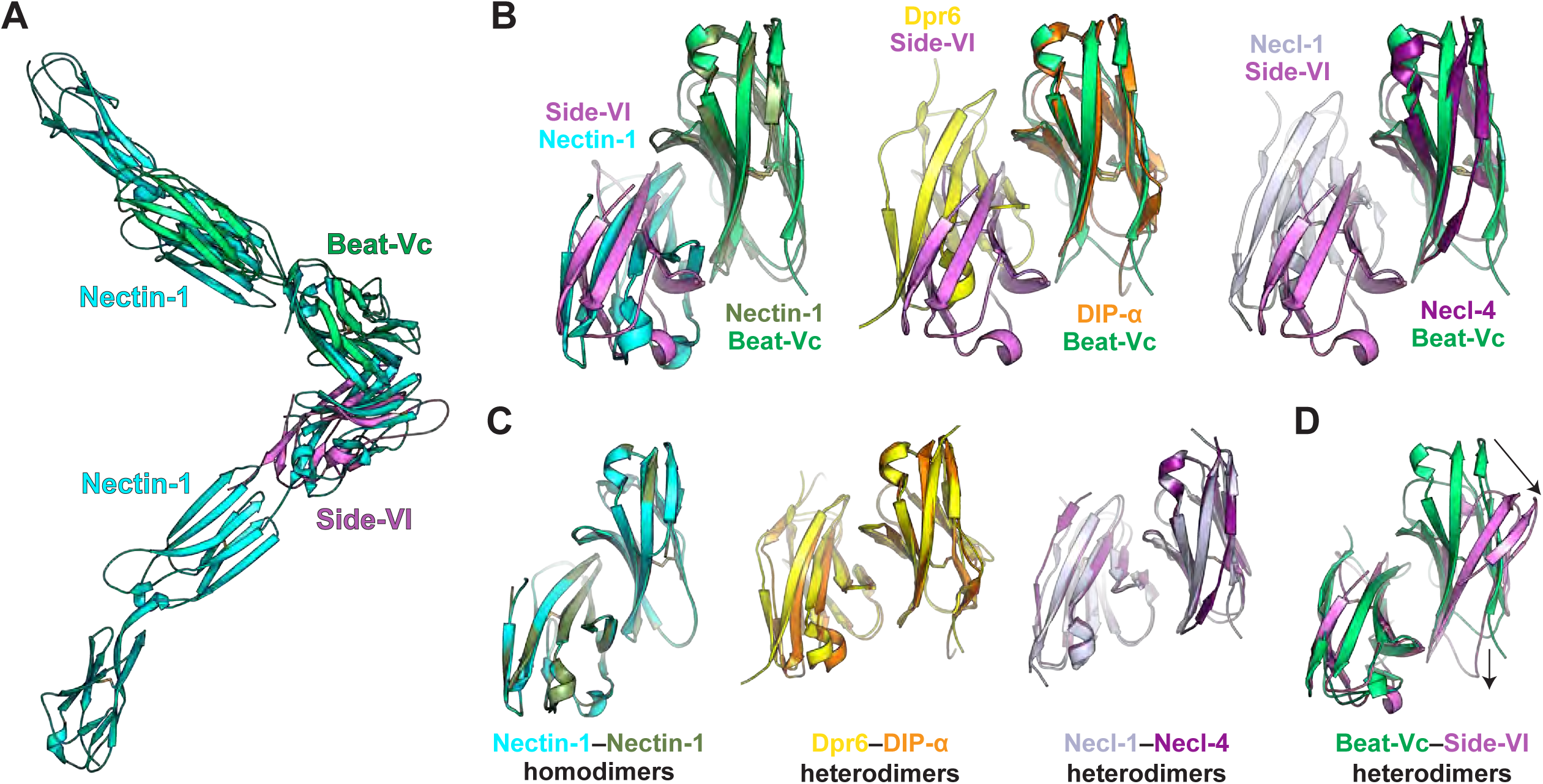
Beat-Vc–Side-VI complex is structurally similar to Nectin and Dpr/DIP complexes. **A.** Overlay of the Beat-Vc–Side-VI complex with the Nectin-1 ectodomain homodimer (PDB ID: 3ALP). Sensograms and binding isotherms for SPR experiments testing the interaction of Beat-Ia and Side. **B.** Overlay of the Beat-Vc–Side IG1-IG1 complex with IG1-IG1 complexes from Nectin-1 homodimer, Dpr6–DIP-α heterodimer, and Necl-1–Necl-4 heterodimer. For the superpositions, Beat-Vc IG1 was aligned with one subunit to highlight differences in the topology of the complexes. **C.** Nectin-1 homodimer, Dpr6-DIP-α and Necl-1–Necl-4 heterodimers are two-fold symmetric IG1-IG1 dimers, as seen when a second copy of the complex is superimposed after rotating the complex, matching different subunits. **D.** Beat-Vc–Side-VI complex does not exhibit two-fold rotational symmetry. When a second of the complex is superimposed on the first by aligning a Beat-Vc and Side-VI, the other monomers do not align well with each other, unlike the Nectin/Necl and Dpr/DIP complexes above.

An invariant feature of Nectin/SynCAM and Dpr–DIP complexes is the strong (pseudo)symmetry of the IG1-IG1 complexes, as seen in the Nectin-1 homodimer and the Dpr6–DIP-α heterodimer (Figure 3C) when the complexes overlay near perfectly with themselves when rotated 180°. Rotating the Beat-Side complex by overlaying Side-VI IG1 domain on Beat-Vc IG1 domain, however, showed that the Beat-Side IG-IG complexes do not have this two-fold symmetry (Figure 3D).

Last, we compared the various Beat IG1+2 models we observed in our crystal structures. While we observed no significant differences between the two copies of the Beat-Vc–Side-VI complex (Figure S3A), our Beat-Vb and Beat-Vc IG1+2 structures show significant movement around the IG1-IG2 domain boundary (Figure S3A), suggesting flexibility or variability in or between Beat ectodomains.

### Chemical features of the Beat–Side interface and the underlying rules of specificity

Utilizing this first structure of a Beat–Side complex, we studied the molecular interface between Beats and Sides. The interface is significantly conserved, as many interface residues, shown with magenta and green boxes in Figure 4A, are invariant across all *Drosophila* Beat and Side sequences, and most other positions are strongly conserved. One feature, shared closely with Nectin/SynCAM complexes and not with Dprs and DIPs, is charged residues throughout the interface, and especially at the core. Beat-Vc and Side-VI surfaces show excellent charge complementary at the interface (Figures 4C,D). In fact, the mutation of most of these amino acids to alanine results in a full or near-complete loss of Beat-Vc–Side-VI binding, as tested in Figure 2F. The amino acid position following the *F* strand cysteine, which is at the center of the interface, is always a Glu in Beats, and an Arg in Sides; this charge-based complementarity is also observed in Nectins, and is an underlying restraint for why certain Nectin complexes, such as the heterodimeric Nectin-1–Nectin-3 complex, can form while others do not (box in Figure 4D).^19^ Overall, these observations demonstrate that the polar interactions at the core of the interface are crucial for Beat-Side binding.

**Figure 4.**
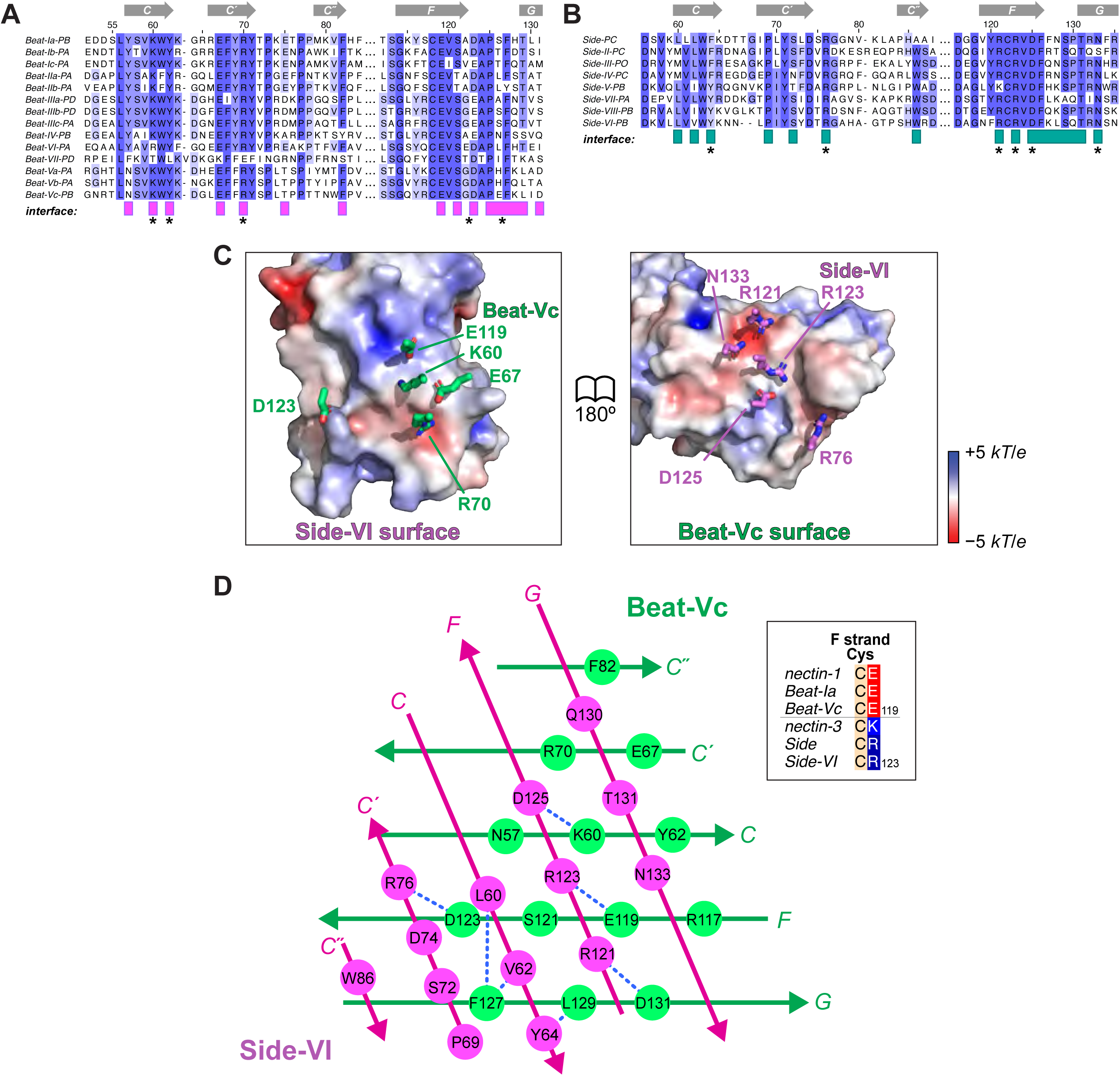
Sequence and structural features of the Beat–Side interface. **A,B.** Sequence alignments of *D. melanogaster* Beats and Sides around amino acids mediating their complex. Sequence numbering above the alignment reflect that of Beat-Vc (A) and Side-VI (B). Residues found within 4 Å of the other subunit in the Beat-Vc–Side-VI structure are labeled with colored boxes under the sequence alignments. Positions with mutations shown to disrupt Beat-Vc–Side-VI binding in Figure 2F are labeled with an asterisk below the alignments. **C.** Side-VI and Beat-Vc surfaces show charge complementarity in an open book view of the interface. **D.** Schematic view of interactions between the beta-sheets of Beat-Vc and Side-VI, highlighting favorable non-covalent interactions (blue dashes), including both polar and non-polar. The amino acid position following the F-strand Cys is highlighted to be a Glu in all Beats and Arg/Lys in all Sides (see inset); this amino acid is also a major determinant of Nectin–Nectin specificity, as observed in the mouse Nectin-1–Nectin-3 complex.

It should be noted that hydrophobic interactions are also present at the Beat-Side interface, specifically where the Beat *G* strand meets the Side *C* strand (Figure 4D). These positions (such as Beat-Vc F127) are also invariantly hydrophobic across all Beats and Sides. Finally, as the resolution of diffraction data is limited to 3.9 Å, the exact side chain positions are often not observed in our electron density maps. In addition, we only have one Beat-Side complex structure. These limitations impede our ability to determine why certain Beats and Sides complex while others do not.

### Solution Behavior of Beat and Side ectodomains

The first functional roles for Beats and Sides were described for the founding members of the two families, Beat-Ia and Side, in motor neuron branching and neuromuscular junction (NMJ) formation in embryos. *Beat-Ia* and *side* phenotypes in NMJ formation are also strongly penetrant, unlike phenotypes for other Beat and Side genes.^17,20^ Therefore, we next characterized Beat-Ia–Side complex formation. We purified Beat-Ia and Side ectodomains and observed that Side runs on size-exclusion chromatography columns at elution volumes matching that of a dimeric size or higher. The reconstituted complex similarly runs at an elution volume expected of a heterotetramer or larger (Figure 5A). Since guidance cues are often oligomers and cause oligomerization of their receptors, we performed multi-angle light scattering (MALS) experiments for the Side ectodomain to measure its molar mass (Figure 5B). Surprisingly, we observed an average mass of 93 kDa, close to the expected monomer of 84 kDa (including predicted N-glycans). Small-angle x-ray scattering (SAXS) showed that Side ectodomains are highly elongated in shape and flexible, which would explain why Side appears as a larger molecule on SEC runs than expected (Figure 5C and S4). Similarly, the complex molar mass (measured: 148 kDa) is closer to that of a heterodimer (expected: 123 kDa) than a heterotetramer (Figures 5D,E and S4). However, based on these results, we cannot rule out Sides oligomerizing weakly.

**Figure 5.**
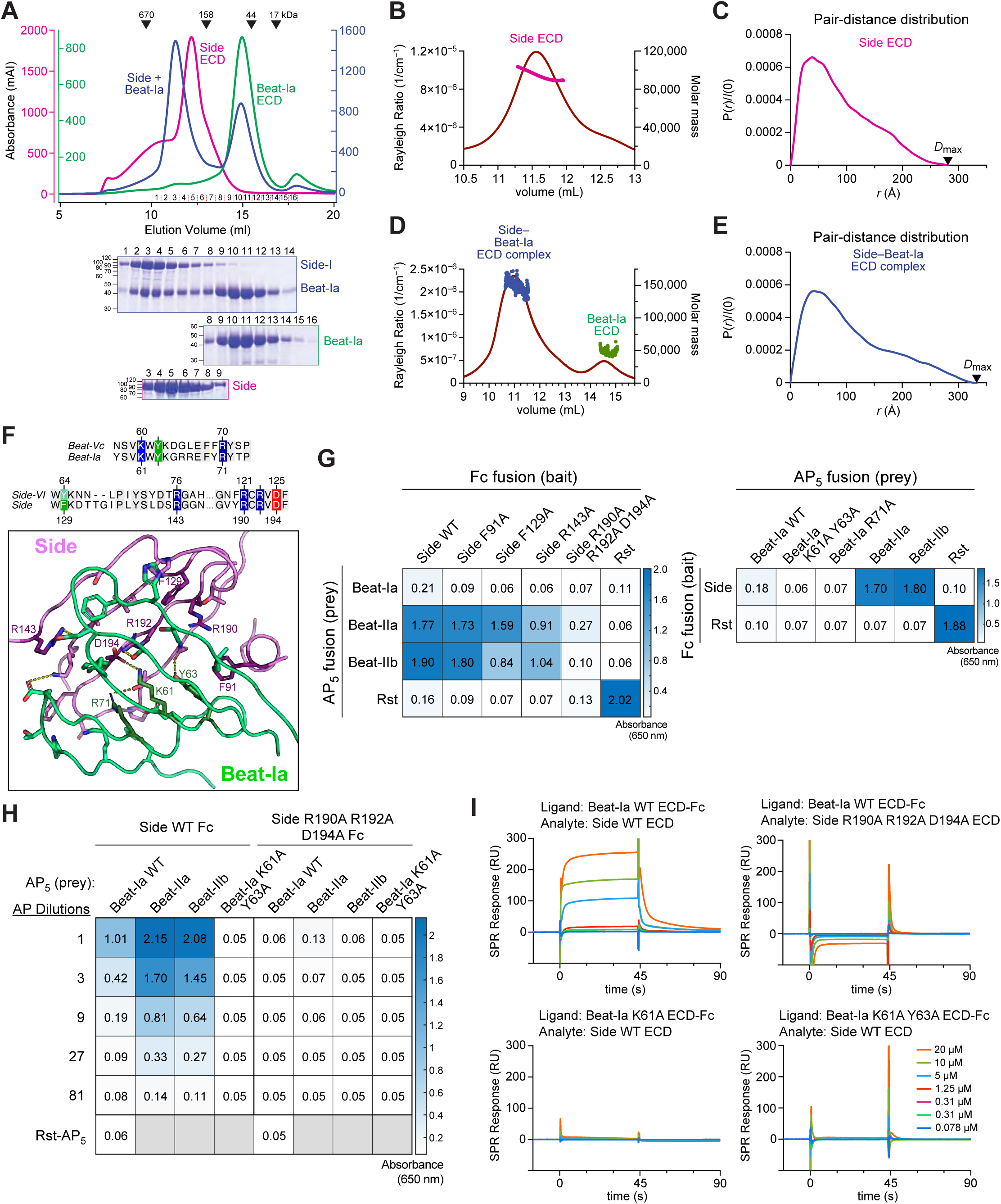
Biophysical characterization and engineering of the Beat-Ia–Side complex. **A.** Size-exclusion chromatography runs for ectodomains of Beat-Ia, Side, and their complex. The column used is a Superdex 200 Increase 10/300, and absorbance is reported from a UV cell with a 0.5 cm pathlength. Elution volumes for size standards are reported with triangles above. Coomassie-stained SDS-polyacrylamide gels for the separate and mixed runs are placed under the chromatograms. **B,D.** Multi-angle light scattering for Side ECD (B) and Beat-Ia (ECD (D), showing a number-average molar mass (*M*_n_) of 93.4 kDa for Side ECD (magenta), 147.6 kDa for Side–Beat-Ia ECD complex (blue) and 46.9 kDa for Beat-Ia ECD (green). **C,E.** Pair-distance distribution plots for Side ECD (C) and Side–Beat-Ia ECD (E), showing highly extended shapes for both and *D*_max_ values of 281 Å and 333 Å, respectively. **F.** Homology modeling of the Beat-Ia–Side complex structure. Residues mutated in Beat-Ia and Side to break their interaction are highlighted in the sequence alignment above, and with darker shades of magenta and green in the structural model below. **G.** ECIA results for the binding of engineered Side mutants to Beat-Ia, IIa and IIb (left), and of engineered Beat-Ia mutants to Side (right). **H.** ECIA with the selected mutations, Beat-Ia K61-Y63A and Side R190A-R192A-D194A, where binding to prey (Beat) was titrated down in a series of dilutions. Absorbance values for the dilution series are plotted in Figure S5C. **I.** SPR sensorgrams show that while wild-type Beat-Ia-Fc on an SPR chip interacts wild-type, the engineered mutants show no binding.

### The Beat-Side binding interface is conserved in the Beat-Ia–Side complex

The structural information we have is currently limited to the Beat-Vc–Side-VI pair. To test if the binding interface is conserved between different Beat-Side pairs, we modeled the Beat-Ia–Side IG1-IG1 complex using our Beat-Vc–Side-VI structure, and predicted that conserved interactions occur at the Beat-Ia–Side interface (Figure 5F). When we mutated Beat-Ia and Side with amino acid substitutions that disrupted the Beat-Vc–Side-VI complex, we observed loss of binding of Side with Beat-Ia, Beat-IIa and Beat-IIb using ECIA (Figures 5G,H and S5). We further confirmed using surface plasmon resonance (SPR) experiments that no binding could be observed for Side ectodomain with the triple R190A-R192A-D194A mutation using a Beat-Ia-captured SPR chip surface (Figure 5I). Similarly, wild-type Side ectodomains elicited no binding response to Beat-Ia with the K61A-Y63A double mutation. For functional studies in the rest of this manuscript, we named these two mutants Side^ΔB^ and Beat-Ia^ΔS^, as Side and Beat-Ia variants with no Beat or Side binding, respectively.

### Beat-Ia–Side interactions are required for motor neuron-muscle connectivity

Previous studies demonstrated a role for Beat-Ia and Side in the developing *Drosophila* neuromuscular system.^5,7,18,21,22^ Therefore, we used this system as an *in vivo* model to validate the Beat-Ia and Side binding interface we identified. *Drosophila* larvae have highly stereotyped hemisegments comprised of 30 muscles innervated by ∼33 motor neurons with nearly invariant innervation patterns (Figure 6A). Using CRISPR, we generated the triple R190A-R192A-D194A *side* mutant (hereafter referred to as *side^ΔB^*), which we showed abrogated binding to Beat-Ia (Figures 5G-I). Third instar homozygous *side^ΔB^*larvae showed reductions in motor neuron innervation of four ventral longitudinal (VL) muscles, m6, 7, 12, and 13, compared to heterozygous control larvae (Figure 6B-D). To test if disrupting the Beat-Ia-Side interface mimics the *side* null mutant, we generated transallelic combinations of *side^ΔB^* with two null alleles, *side^I156^* and *side^C1^*^37^.^6,7,18,23^ While these transallelic larvae (*side^ΔB^*/*side^I^*^1563^ and *side^ΔB^*/*side^C1^*^37^) showed less innervation of VL muscles than homozygous *side^ΔB^* larvae, *side* null larvae (*side^I^*^1563^/*side^C1^*^37^) were even more penetrant (Figure 6D, S6). In both *side^ΔB^*/*side^I^*^1563^ and *side^ΔB^*/*side^C1^*^37^ larvae, we observed innervation of VL muscles at improper locations, suggesting either incorrect motor axon trajectories or ectopic innervation by neighboring motor neurons (Figure S6). Overall, these results suggest that the Beat-Ia–Side interaction interface determined by structural and biochemical analyses is critical for Side function *in vivo*.

**Figure 6.**
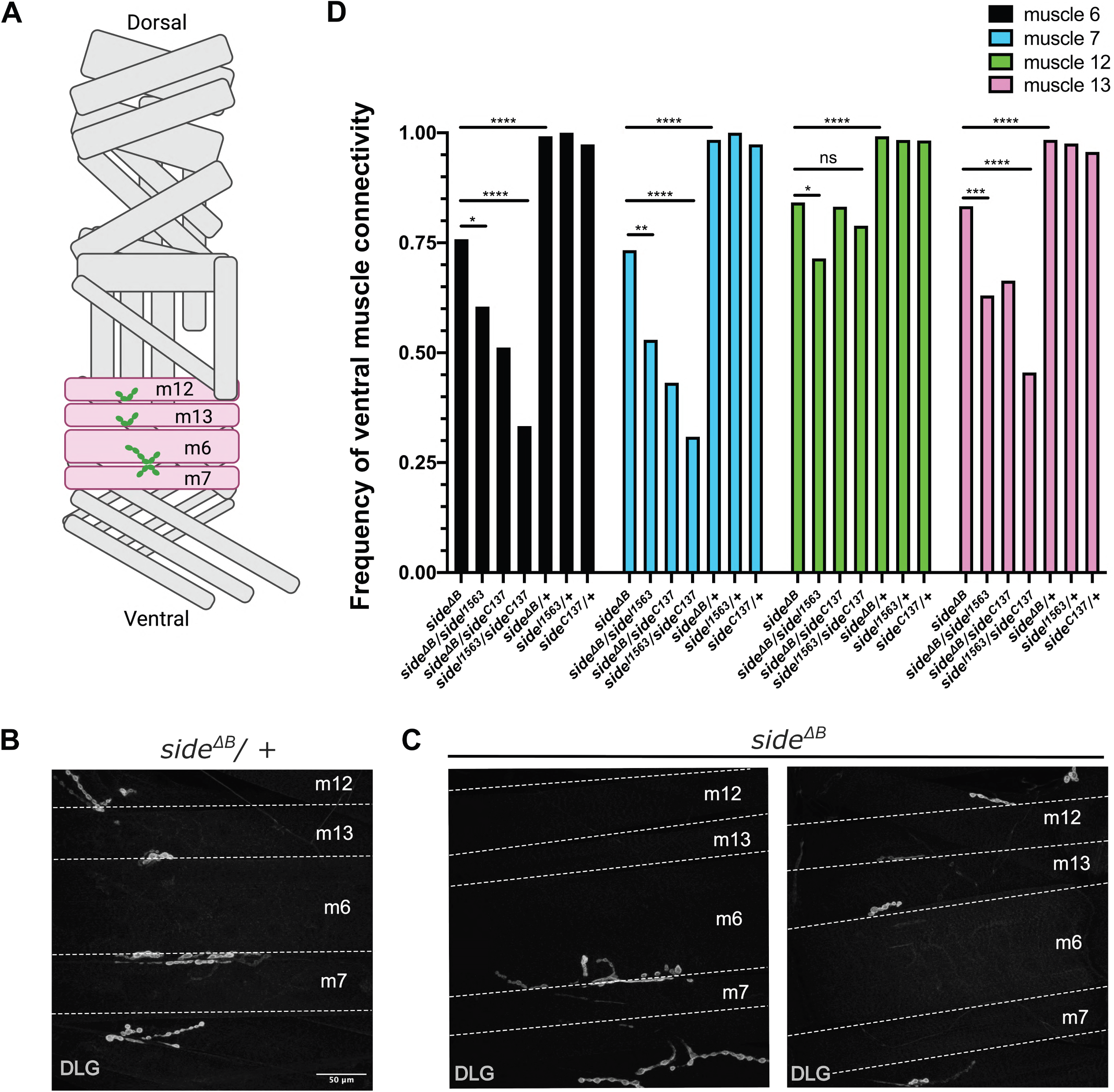
*side^ΔB^* larvae showed reduced connectivity on ventral muscles. **A.** Schematic of a third instar larval body wall hemisegment depicting the wild-type innervation pattern on ventral longitudinal muscles, highlighted in pink. **B,C.** Representative images of the innervation patterns in (B) heterozygous control *side^ΔB^/+* or (C) homozygous *side^ΔB^*. Postsynaptic membranes on muscles were labeled with anti-DLG (white). **D.** Quantification of connectivity frequency on ventral longitudinal muscles. (Fisher’s Exact test, * p ≤0.05, ** p ≤0.01, *** p ≤0.001, **** p <0.0001, ns >0.05; Sample size: *side^ΔB^*, n = 16; *side^I1563^/side^ΔB^*, n = 15; *side^C137^/side^ΔB^*, n = 16; *side^C137^/side^I1563^*, n = 16; *side^ΔB^/+*, n = 16; *side^I1563^/+*, n = 16; *side^C137^*/+, n = 15)

### Beat-Ia–Side interaction is required for side gain-of-function phenotype

Side is dynamically expressed on different cells in the nervous system and muscle fields to guide motor neuron axons migrating toward the appropriate muscle targets. However, constitutive overexpression of Side in developing muscles leads to a gain-of-function phenotype that attracts the outgrowing axons prematurely to muscle fields and inhibits their migration and innervation of dorsal muscles.^7,18^ To test if Beat-Ia–Side interactions are required for this gain-of-function phenotype, we generated transgenic UAS-Side and UAS-Side^ΔB^ lines. We also generated corresponding myc-tagged variants for both wild-type and mutant Side. When overexpressed in muscles, myc-tagged Side and Side^ΔB^ localized to the muscle surface and aggregated around the motor axon terminals (Figure S7A-B). Overexpression of Side or Side-myc led to a significant loss of innervation on the most dorsal muscles, m1 and 9. However, overexpression of Side^ΔB^ or Side^ΔB^-myc revealed an innervation frequency similar to wild type (Figure 7). Overexpression of Side or Side^ΔB^ did not affect innervation of m2 or m10 or other more ventral muscles (Figure S7C-D). This gain-of-function phenotype was reported to be suppressed by coexpression of Beat-Ia in muscles.^7^ We generated transgenic lines for UAS-Beat-Ia and UAS-Beat-Ia^ΔS^, which carries the K61A-Y63A double mutations shown to disrupt Beat-Ia binding to Side (Figure 5G-I). Coexpression of wild type UAS-Beat-Ia with UAS-Side-myc alleviated the loss of connectivity on m9 shown in muscle overexpression of UAS-Side-myc alone. However, muscle coexpression of UAS-Beat-Ia^ΔS^ did not suppress this gain-of-function phenotype (Figure 7E). These results suggest that the Beat-Ia–Side interface drives the Side gain-of-function phenotype and bolster the model whereby Beat-Ia–Side interactions instruct motor axon guidance and assembly of the neuromuscular circuit.

**Figure 7.**
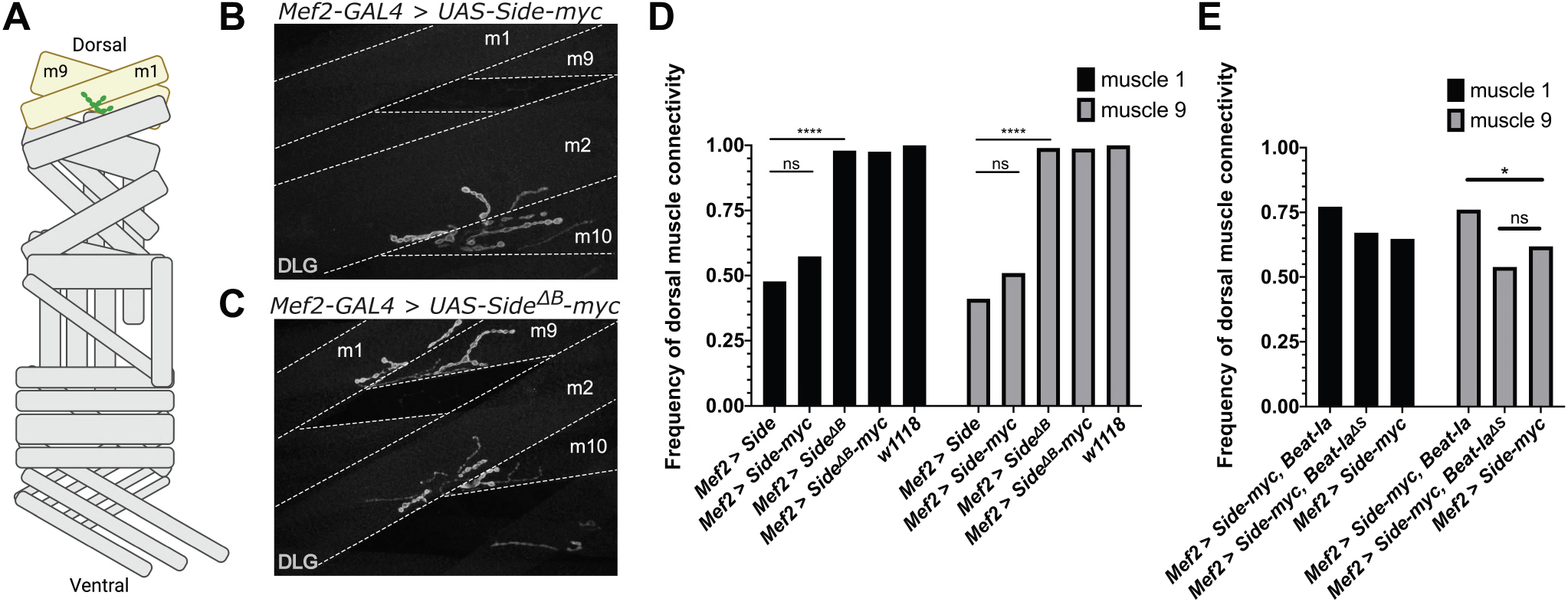
Loss of Side-Beat-Ia interaction rescues the gain-of-function Side innervation phenotype. **A.** Schematic of a third instar larval body wall hemisegment depicting the wild-type innervation pattern on dorsal muscles 1 and 9, highlighted in beige. **B,C.** Representative images of innervation on dorsal muscles m1 and m9 in third instar larvae with (B) wild type Side muscle overexpression or (C) Side*^ΔB^* muscle overexpression. Postsynaptic membranes on muscles were labeled with anti-DLG (white). **D.** Quantification of connectivity frequency on muscle 1 and 9. (Fisher’s Exact test, * p ≤ 0.05, ** p ≤ 0.01, *** p ≤ 0.001, **** p < 0.0001, ns > 0.05; Sample size: *Mef2 > Side*, n = 22; *Mef2 > Side-myc*, n = 24; *Mef2 > Side^ΔB^*, n = 22; *Mef2 > Side^ΔB^-myc*, n = 24; *w^1118^*, n = 18) **E.** Quantification of connectivity frequency on muscle 1 and 9 upon coexpression of Side-myc and Beat-Ia variants with or without double mutations. (Fisher’s Exact test, * p ≤ 0.05, ns > 0.05; Sample size: *Mef2 > Side-myc, Beat-Ia*, n = 23; *Mef2 > Side-myc, Beat-Ia^ΔS^*, n = 21; *Mef2 > Side-myc*, n = 24)

## DISCUSSION

It is not fully resolved how the wide collection of cell surface molecules and signaling receptors presented on neuronal surfaces instructs circuit assembly. In the *Drosophila* neuromuscular system, Beaten path and Sidestep were among the first cell surface molecules implicated in motor neuron axon branching and innervation, acting as an attractive guidance receptor-cue pair. Since then, Beats and Sides have been implicated in the development of visual circuits and the adult neuromuscular circuits in *Drosophila*. However, the biochemical characterization of their interaction, and their closely related paralogs in the Beat and Side families, have remained limited.

Here, we show that the Beat–Side interaction is mediated by the IG1 domains of both proteins, as also shown recently by Heymann et al.,^18^ and report the first structure of a Beat–Side complex. We validate the structure by a mutational analysis of our observed Beat-Vc–Side-VI complex interface and the homology-inferred Beat-Ia–Side interface. The interaction is mediated by the *CCʹCʺFG* sheet of both proteins, as seen before in IG-IG cell surface adhesion complexes prevalent in the nervous system, most importantly Nectin and SynCAM/Necl, and Dpr and DIP families. The prevalence of this mode of interaction in neuronal IgSF receptors suggests a bilaterian ancestor to the aforementioned protein families, often associated with neuronal wiring and synapses; however, evolutionary relationships between these families are difficult to establish, especially in the case of Beats and Sides, and we cannot exclude the possibility that some of these similarities are the result of convergent evolution.

Using structural comparisons, we have observed a stronger similarity of the Beat–Side complex to the complexes formed by Nectins and SynCAM/Necls. These complexes share a mostly polar interface at their core, unlike Dpr–DIP complexes. Here we highlight and suggest specific amino acid positions that may be (1) important for Beat–Side specificity, and (2) should be targeted to break Beat–Side complexes in future *in vitro* and *in vivo* studies. We expect that our structural and biochemical data serves as a groundwork for future studies of Beat–Side and wider IG-IG specificity in IgSF cell adhesion molecules.

Axon guidance cues often oligomerize their receptors. Our experimental observations suggest that the Side ectodomain is mostly monomeric, and that the Beat-Ia–Side complex may only weakly oligomerize, if at all. Therefore, it remains unclear how Beats are activated by Sides and if ligand-mediated oligomerization plays any role in the process. However, it is still possible that Sides oligomerize via their cytoplasmic domains, e.g., through PDZ domain interactions, with scaffolding proteins intracellularly, or by interactions with co-receptors, which may result in Beat clustering and activation.^17^

Our structural and biochemical analyses revealed that site-specific mutations in Side effectively disrupt its interaction with Beat-Ia. We reasoned that introducing these mutations into *Drosophila* would mimic muscle innervation phenotypes reported for *side* null mutants. While we did observe defects in muscle innervation, the *side^ΔB^* phenotypes were not as penetrant as null mutants. One possibility is that Side may interact with other cell surface proteins, in addition to Beat-Ia, to instruct innervation of muscles. While the Side mutations specifically abrogate binding to Beat-Ia, the interaction between Side and other proteins could be preserved and thus retain some of the Side function. A recent study showed that Side-IV uses the cell surface protein Kirre as a co-receptor in instructing specific synaptic connectivity in the adult *Drosophila* visual system.^17^ Whether Side or Beat-Ia interact with other cell surface proteins to regulate larval neuromuscular system development is unknown. In addition, we observed that on *side^ΔB^*muscles that retained motor neuron innervation, the axon sometimes took alternative trajectories and innervated at inappropriate locations. Therefore, we cannot rule out the possibility that *side^ΔB^* animals showed defects in a comparable number of muscles to *side* null mutants because these milder phenotypes were not quantified in this study.

We also noticed that the loss of dorsal muscle innervation upon pan-muscle overexpression of wild-type Side was milder compared to previous studies.^7,18^ In these prior studies, ectopic expression of the wild type UAS-Side driven by Mef2-GAL4 led to a significant loss of motor neuron connectivity on dorsal muscles m2 and 10, which was not observed in our study. One possible explanation is that the expression levels of the Side transgenes were different due to genetic background, insertion site, and/or construct design. However, the innervation defect at muscles m1 and m9 suggests that our transgenic lines function properly. Future experiments are needed to determine if expression levels are critical for Beats and Sides, as reported for other CSPs.^24–26^

## Acknowledgments

This work was funded by National Institutes of Health (NIH) grants R01 NS139060 (to E.Ö and R.A.C.), R01 NS123439 (to R.A.C), T32 GM138826 (to J.M.P.), T32 GM144292 and F31 NS136004 (to R.L), and R01 NS028182 (to E.Ö). We thank the Bloomington Drosophila Stock Center (NIH P40OD018537) for fly lines. We thank Hermann Aberle for sharing the *side^I1563^*and *side^C137^* lines, Maxwell Watkins for advice with SAXS data analysis and K. Christopher Garcia for reagents and advice. We thank x-ray beam lines and staff at the Advanced Photon Source (APS) sectors GM/CA, NECAT and BioCAT, and at the Stanford Synchrotron Radiation Lightsource (SSRL) for assistance to collect data. GM/CA at the APS has been funded by the National Cancer Institute (ACB-12002) and the National Institute of General Medical Sciences (NIGMS) (AGM-12006, P30GM138396). The Northeastern Collaborative Access Team (NECAT) beamlines are funded by the NIGMS from the NIH (P30 GM124165). This research used resources of the APS, a U.S. Department of Energy (DOE) Office of Science User Facility operated for the DOE Office of Science by Argonne National Laboratory under Contract No. DE-AC02-06CH11357. The SSRL at the SLAC National Accelerator Laboratory is supported by the U.S. DOE, Office of Science, Office of Basic Energy Sciences under Contract No. DE-AC02-76SF00515. BioCAT was supported by grant P30 GM138395 from the NIGMS of the NIH. The SSRL Structural Molecular Biology Program is supported by the DOE Office of Biological and Environmental Research, and by the NIH, NIGMS (P30GM133894). The monoclonal 4F3 was developed by C. Goodman and obtained from the Developmental Studies Hybridoma Bank, which was created by the NICHD of the NIH and maintained at the University of Iowa, Department of Biology.

## Author Contributions

A.M.O and E.Ö. determined crystal structures. J.M.P. and A.B.C. engineered proteins and performed various binding experiments. A.B.C. and E.Ö. performed x-ray and light scattering experiments. A.M.O, J.M.P and A.B.C. expressed and purified proteins. R.Z. and J.A. generated mutant animals. R.Z. performed imaging and in vivo analysis. E.Ö., J.M.P., R.Z. and A.B.C. wrote the manuscript with input and review from all authors.

## METHODS

### Extracellular Interactome Assay (ECIA)

Secreted Fc-tagged (bait) and AP5-tagged (prey) proteins for ECIA were expressed in *Drosophila* S2 cells using transient transfections with TransIT-Insect (Mirus Bio) in Schneider’s medium (Thermo Fisher, cat. no. 21720001) supplemented with 10% Fetal Bovine Serum, 50 units/ml Penicillin, 50 µg/ml Streptomycin, and 2 mM L-Glutamine.^27^ Conditioned media were run on SDS-PAGE gels for western blotting with a mouse anti-His Tag Antibody [iFluor 488] (GenScript, A01800) to confirm protein expression and to normalize protein concentrations between wild-type and mutant forms of the same protein, allowing for quantitative comparisons.

For the assay, Protein A-coated 96-well plate wells were washed with Phosphate-Buffered Saline (PBS) with 0.1% Tween-20 (PBST). To capture bait, 100 µl of medium with secreted Fc-tagged bait was added to each well for overnight incubation while shaking at 4°C. Wells were blocked with PBS with 1% BSA at room temperature for 3 hours while shaking. Wells were then washed with PBS with 1 mM CaCl_2_, 1 mM MgCl_2_, and 0.1% BSA, and incubated with 100 µl of medium with secreted AP-tagged prey at room temperature for 3 hours while shaking. After a final wash with PBS with 1 mM CaCl_2_, 1 mM MgCl_2_, and 0.1% BSA, 100 µl of BluePhos Phosphatase Substrate (Seracare, cat. no. 5120-0059) was added to each well. To assess bait-prey binding, absorbance at 650 nm was measured with a BioTek Synergy H1 Plate Reader.

### Protein Expression and Purification for Size Exclusion Chromatography and Surface Plasmon Resonance

For biophysical studies, proteins were expressed using the baculoviral expression system and the lepidopteran High Five cell line in Insect-XPRESS medium (Lonza, BELN12-730Q).

Baculoviruses were produced using homologous recombination in insect cells by co-transfection with linearized baculoviral DNA (Expression Systems, 91-002) and the transfer vector pAcGP67A (BD Biosciences), which carries an N-terminal gp64 signal peptide for secretion. All proteins were tagged C-terminally with a hexahistidine tag for purification.

Proteins were purified first with Ni-NTA metal-affinity chromatography (QIAGEN, cat. no. 30210) from conditioned media, followed by SEC with Superdex 200 Increase 10/300 columns (GE Healthcare, cat. no. 28-9909-44). We observed that full ectodomain Beat and Side constructs often aggregated or were completely lost over SEC columns when physiological salt was used. We used HEPES-buffered saline (HBS) with 10 mM HEPES, pH 7.2 and 150 mM NaCl as the base buffer, but had to modify it to include 500 mM NaCl and/or supplement with 10% Glycerol on occasion to ensure lack of aggregation, solubility and stability (see next section for details). Specifically, Beat-Ia ectodomain required 10 mM HEPES, pH 7.2 with 500 mM NaCl and 10% Glycerol during purification, which was used during the reconstitution SEC runs for the Beat-Ia– Side ectodomain complex, shown in Figure 5a.

### Protein Crystallization and Structure Determination by X-ray Crystallography

Beat-Vb IG domains 1+2 were expressed with a C-terminal hexahistidine tag and purified in HBS. 10 mg/ml protein was crystallized using sitting-drop vapor diffusion in 4.8% (w/v) Polyethylene glycol (PEG) 3,350, 22% (v/v) PEG 200 and 0.1 M MES pH 6.0 at 21°C. Crystals were cryo-protected in 5% PEG 3,350, 35% PEG 200 and 0.1 M MES pH 6.0 and vitrified in liquid nitrogen. Diffraction data was processed using XDS.^28^ Phasing was done by molecular replacement in PHASER (version 2.8.3)^29^ using an Alphafold 2 model obtained via the Colabfold implementation.^30,31^

Side-IV (CG14372) IG1 domain was expressed with an N-terminal hexahistidine tag, followed by a TEV protease cleavage site, in B834(DE3) *E. coli* cytoplasm at 37°C to generate misfolded inclusion bodies. For folding of Side family IG1 domains, see next section. Refolded and purified tagless Side-IV IG1 was crystallized at 7.1 mg/ml concentration in HBS using 0.2 M Ammonium sulfate, 15% (w/v) PEG 4,000, 17% Glycerol as the crystallant in sitting-drop vapor diffusion experiments at 21°C. Crystals were cryoprotected in 0.2 M Ammonium sulfate, 20% PEG 4,000, 20% Glycerol. Diffraction data was processed with XDS^28^ and molecular replacement was performed using MrBUMP^32^ with the PDB model 2PND chain A.

Side-VI IG1 domain was expressed with a TEV protease-cleavable N-terminal hexahistidine tag in the BL21(DE3) cytoplasm as inclusion bodies. Refolded and purified tagless Side-VI IG1 was crystallized at 9.8 mg/ml mg/ml concentration in HBS using 1.8 M Sodium acetate trihydrate pH 7.0, 0.1 M Bis-tris propane pH 7.0 as the crystallant in sitting-drop vapor diffusion experiments at 21°C. Crystals were cryoprotected in 1.8 M Sodium acetate trihydrate pH 7.0, 0.1 M Bis-tris propane pH 7.0, 30% Glycerol. Diffraction data was processed with XDS and molecular replacement was performed with MOLREP (version 11.5.04)^33^ using a homology model built from our Side-IV structure (see above) by MODELLER.^34^

Side-VII IG1 domain was expressed using baculoviruses in insect cells, and purified in HBS. 8.5 mg/ml protein was crystallized using the sitting-drop vapor diffusion method in 0.1 M MES pH 6.1 + 25% PEG 4000 + 1% 1,2-Butanediol. These crystals were cryoprotected in 0.1 M MES pH 6.1, 25% PEG4000, 30% Ethylene Glycol, and diffracted anisotropically. Diffraction data was processed with XDS^28^ and STARANISO,^35^ using ellipsoid truncation and anisotropic scaling. Molecular replacement was performed with PHASER (version 2.7.16)^29^ using a homology model built from our Side-IV structure (see above) by MODELLER.^34^

For a complex structure, Beat-Vc IG1+2 domains and Side-VI IG1 domain were produced separately using baculoviruses in insect cells. Beat-Vc IG1+2 had to be purified in HBS with 10% Glycerol, while Side-VI IG1 was stable in HBS. A complex sample was formed by mixing the two proteins at a 1:1 molar ratio and concentrating them down to 9 mg/ml, which crystallized in 1.3 M K/Na tartrate, 0.1 M HEPES pH 8.0 using the sitting-drop vapor diffusion method at 21°C. Crystals were cryoprotected in 2 M Ammonium sulfate, 0.1 M Tris pH 8, 25% Glycerol.

Diffraction was severely anisotropic, where diffraction was limited to 5.0 Å in two of the principal orthogonal axes and 3.1 Å in the third in reciprocal space at an ⟨*I*/*σ*(*I*)⟩ cut-off of 1.2. Diffraction data was processed with XDS^28^ and STARANISO,^35^ using ellipsoid truncation and anisotropic scaling, resulting in a 3.9 Å-resolution dataset. Molecular replacement was performed with PHASER (version 2.8.3)^29^ using our Side-VI IG1 structure and a Beat-Vc IG1+2 Colabfold model.

All molecular replacement and reciprocal space refinement jobs were performed using the graphical user interfaces of the PHENIX and CCP4 packages.^36,37^ Reciprocal space refinement was performed with phenix.refine^38^ and model building in real space was done using Coot (version 0.9.8).^39^ Validation of the chemistry in structural models was performed using tools in Coot and with MOLPROBITY.^40^

The coordinates for the structures and the crystallographic structure factors are deposited at the Protein Data Bank with the following accession codes: 9NP7, 9NQ2, 9NS8, 9NSA, 9NSF.

### Refolding of IG1 domains of Side family proteins

Folding was performed as previously described for SYG-1 IG domains.^41^ BL21(DE3) cells are used to express Side family IG1 domains for 3 hours at 37°C in Terrific Broth. The cell pellet was resuspended in 50 mM Tris pH 8.0, 100 mM NaCl, 1 mM EDTA. 1% Triton X-100 before freezing. Defrosted pellet was infused with protease inhibitors, DNase and 1 mg/ml lysozyme before sonication. Following centrifugation, the insoluble pellet was solubilized in wash buffers followed by centrifugation to purify insoluble inclusion bodies: the wash cycles included three runs 20 mM Tris pH 8, 25 mM EDTA, 0.5% Triton X-100 and three cycles of 20 mM Tris pH 8, 25 mM EDTA.

100 mg of pure inclusion bodies was denatured in 1 ml of 8 M Urea, 0.1 M MES pH 6.0, 1 mM DTT, centrifuged to remove the insoluble material, and refolded by slow dripping at 4°C into 100 ml of refolding buffer: 10 mM Tris pH 8, 400 mM L-Arg-HCl, 0.25% PMSF, 0.03% (w/v) oxidized glutathione, 0.15% (w/v) reduced glutathione with protease inhibitors. The refolded sample was dialyzed three times against 1 L of 10 mM Tris pH 8, 150 mM NaCl. Misfolded material was removed by centrifugation; the soluble fraction was incubated by Ni-NTA agarose resin to capture and purify hexahistidine-tagged protein. The purified protein was incubated by TEV protease to remove the N-terminal tag. Uncut protein was removed by binding to Ni-NTA agarose; the protease-cut protein sample was run over Superdex 200 Increase 10/300 in HBS to complete purification.

### Surface Plasmon Resonance

SPR experiments were conducted on a Biacore T200 (GE Healthcare) at 22°C. To capture Beat-Ia (wild-type and mutant) on Protein A-coated SPR chips (Cytiva, 29127555), we used S2 cell conditioned media with secreted Fc-tagged Beat-Ia (also used as bait in ECIA experiments). Purified, soluble Side was used as mobile-phase analyte. SPR experiments were done using a buffer containing 10 mM HEPES pH 7.2, 500 mM NaCl, 10% glycerol and 0.05% Tween-20. Titration data was processed and analyzed in Biacore evaluation software (version 1.0) and Prism version 10 (Graphpad).

### Small-Angle X-ray Scattering and Multi-Angle Light Scattering

SAXS was performed at BioCAT (beamline 18ID at the Advanced Photon Source, Chicago) with in-line size exclusion chromatography (SEC) and multi-angle light scattering (MALS) and refractive index measurement (RI) (SEC-MALS-SAXS). The samples were loaded on a Superdex 200 Increase 10/300 GL column (Cytiva) run by a 1260 Infinity II HPLC (Agilent Technologies). Side ECD was run at a 0.6 ml/min flow rate in HBS, while the mixed Side ECD+Beat-Ia sample (1:1.3 molar ratio) was run in 10 mM HEPES pH 7.2, 500 mM NaCl, 10% Glycerol. The flow passed through (in order) a MALS detector (DAWN Helios II, Wyatt Technologies), an RI detector (Optilab T-rEX, Wyatt), a fiber coupled UV cell, and a SAXS flow cell. The flow cell consists of a 1.0 mm ID quartz capillary with ∼20 μm walls. A coflowing buffer sheath is used to separate sample from the capillary walls, helping prevent radiation damage. Scattering intensity was recorded using an EIGER2 XE 9M (Dectris) detector and data was reduced using BioXTAS RAW.^42^ Buffer blanks were created by averaging regions flanking the elution peak and subtracted from exposures selected from the elution peak to create the I(q) vs q curves used for subsequent analyses. SAXS data were processed in BioXTAS RAW version 2.3.0,^42^ using tools from the ATSAS package, version 4.0.1^43,44^ and other methods implemented in RAW.^45–47^

MALS data were used to calculate molar mass using ASTRA version 6.1.17 software (Wyatt). For mass analysis of Side ECD, we used refractive index as the concentration method and a dn/dc value of 0.1817, based on predicted N-glycan content. For the Beat-Ia+Side ECD run performed in the high salt+glycerol buffer, we used an extinction coefficient of 1.049 mL/(mg×cm) for the Beat-Ia–Side complex and 1.118 mL/(mg×cm) for the Beat-Ia ECD as the method of concentration measurement. See Table S2 for details.

### Modeling and Visualization of other Beat-Side complexes

Beat-Ia–Side-I complex was modeled using MODELLER v. 10 using our Beat-Vc–Side-VI crystal structure.^48^ The Beat-Ia–Side model was also in general agreement with an *Alphafold-multimer*-predicted complex, obtained via the Colabfold implementation (Alphafold v. 2.3.0, Colabfold v. 1.5.2),^49,31^ with an RMSD of 1.4 Å for 191 over 225 Cα atoms aligned. PyMOL (v. 3.1.1) was used for graphical visualization and analysis of all structures.

### Molecular cloning and generation of Side variants

For the Side^ΔB^ endogenous mutation, an 899bp fragment spanning exon 3 and 4 of Side was synthesized by Azenta, and directionally cloned into the pUC57_Kan_gw_OK2.^50^ This fragment contains gRNA1 and gRNA2 (see below) as well as Left homology arm (LH)_SideΔB (see below), a restriction enzyme cassette containing BbsI, and a RH_Side (see below). This construct was then digested with BbsI and a 3xP3_dsRed cassette was amplified from pHD-ScarlessDsRed (DGRC Stock 1364; https://dgrc.bio.indiana.edu//stock/1364; RRID:DGRC_1364) using BsaI (restriction sequences were designed to overlap for directional cloning). The final plasmid was sent to Bestgene (thebestgene.com) for injection into the nos-Cas9 background. Surviving offspring were screened for the 3xP3_dsRed expression, and balanced. All UAS-Side constructs were PCR amplified using Q5 High-Fidelity DNA polymerase (NEB) from cDNA with the below primers, and two UAS-Beat-Ia constructs were PCR amplified from the UAS-Beat line kindly provided by the Aberle lab.^5,18^ The corresponding fragments were combined into the NotI site of pUASTattB using NEBuilder HIFI Assembly (NEB). These constructs were sent to BestGene (thebestgene.com) for injection and integrated into the VK37 landing site (BDSC 9752).

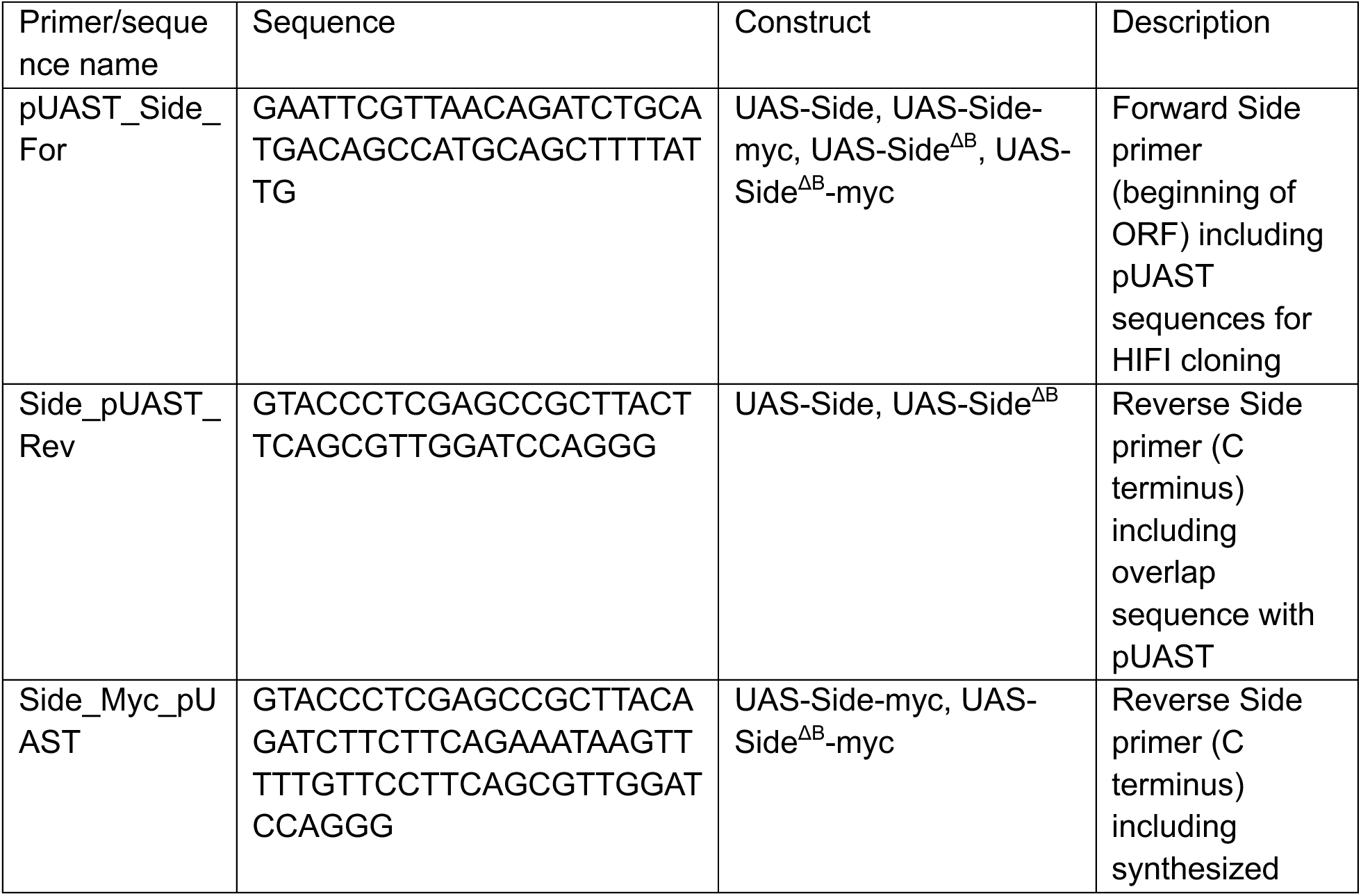
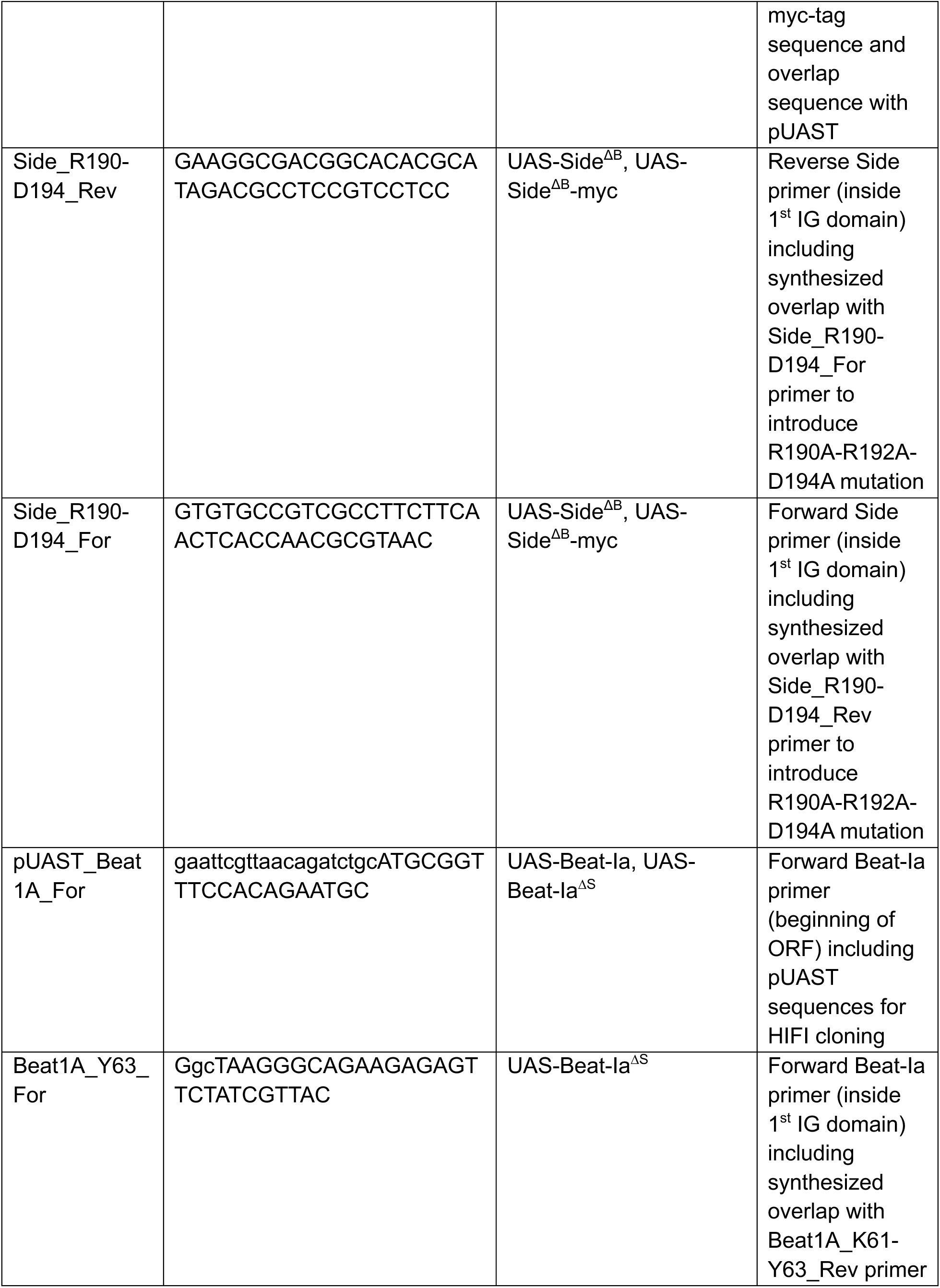
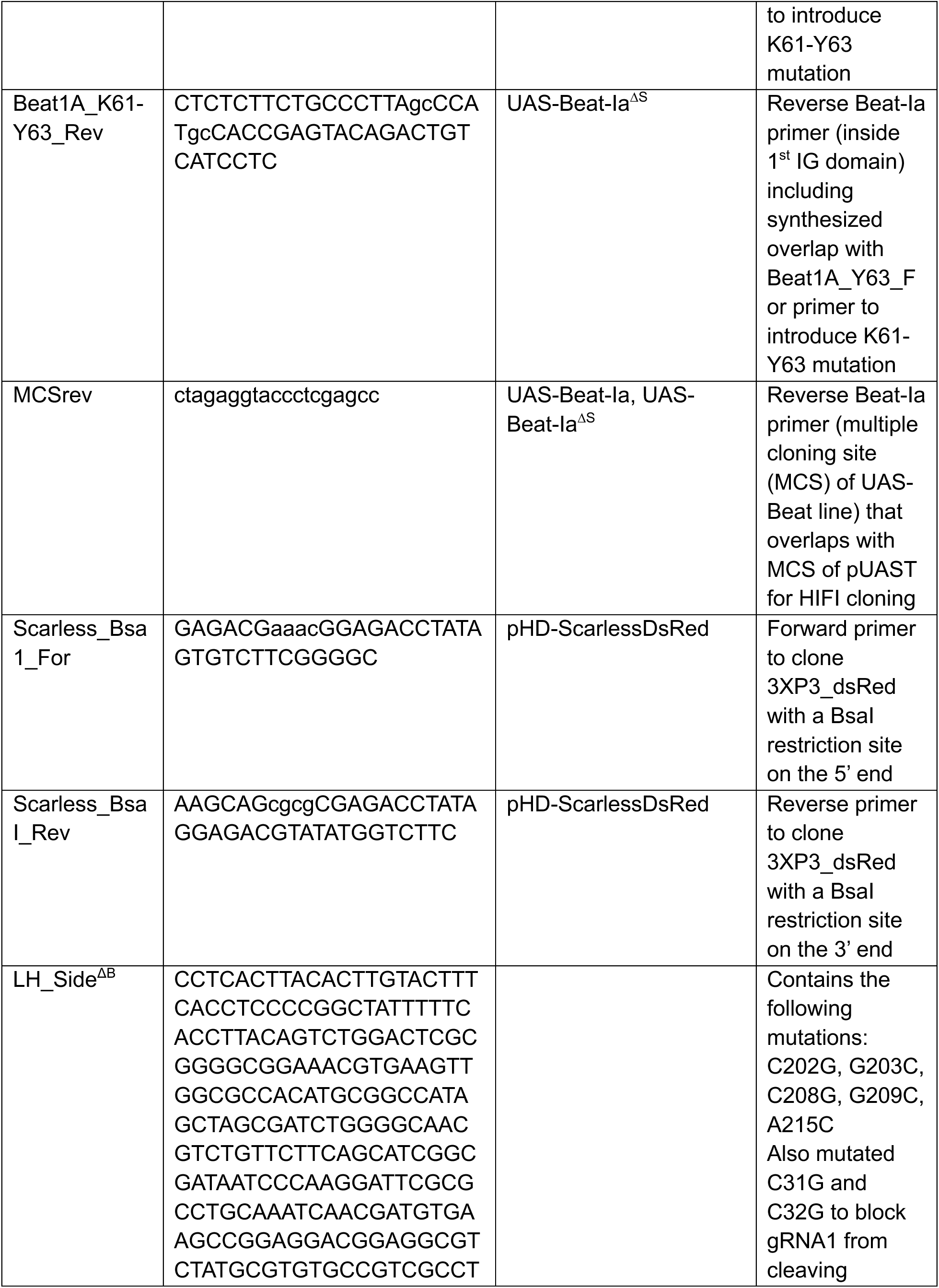
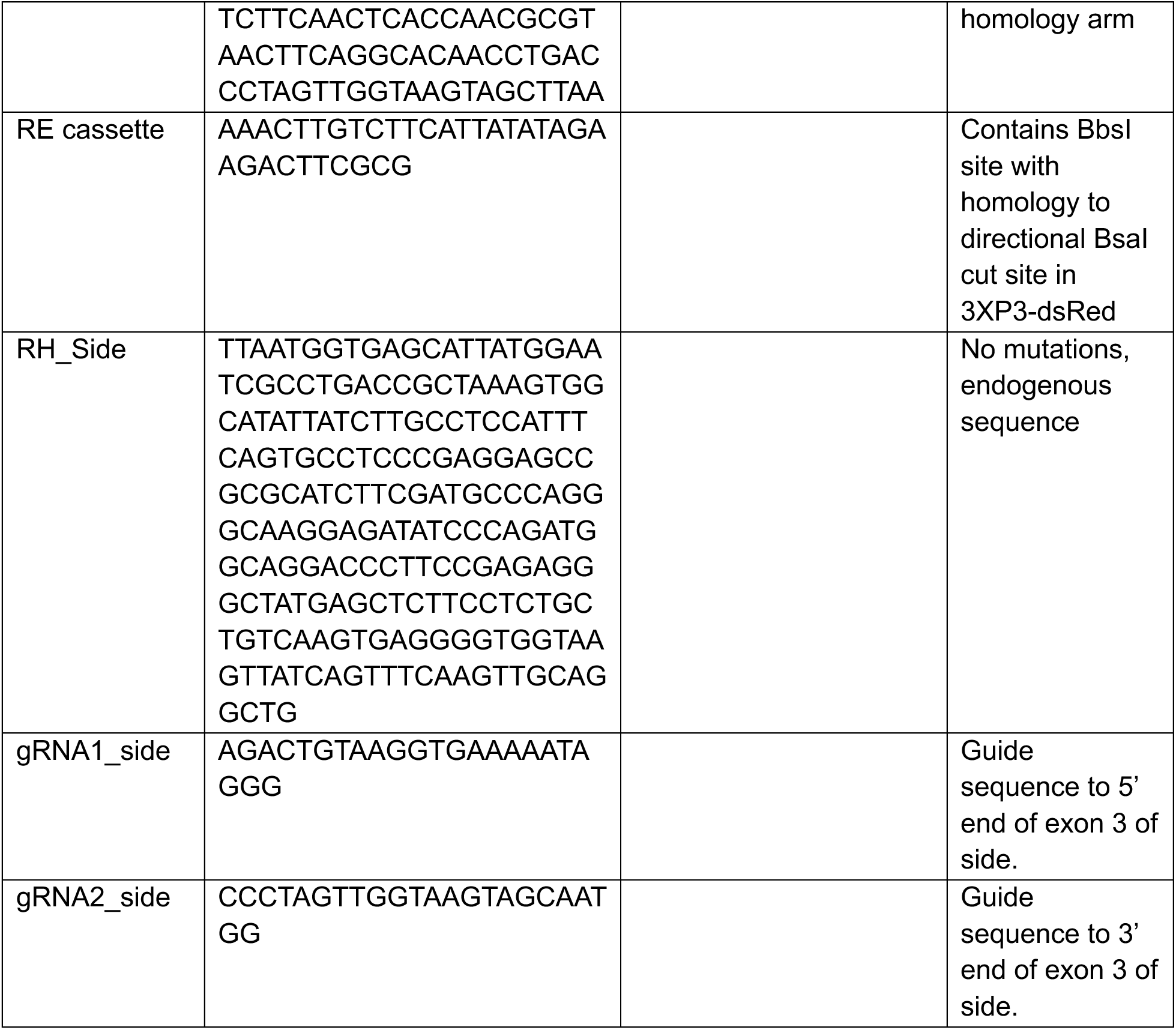

### Drosophila *reagents*

The following lines were used in this study:

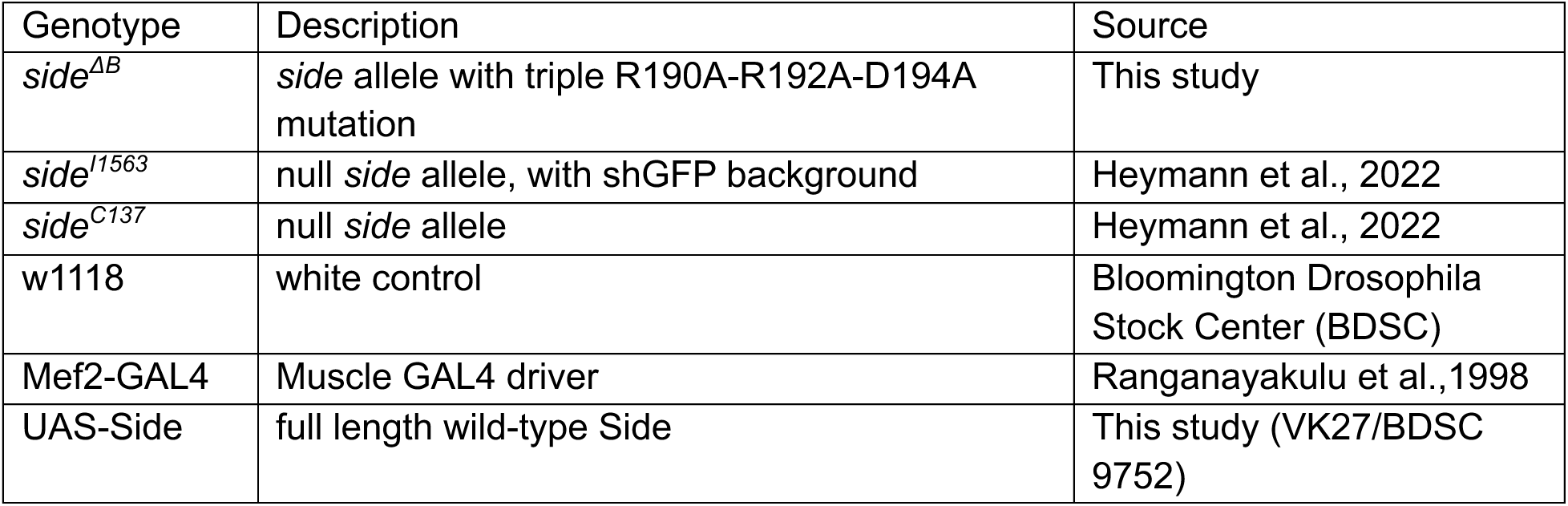
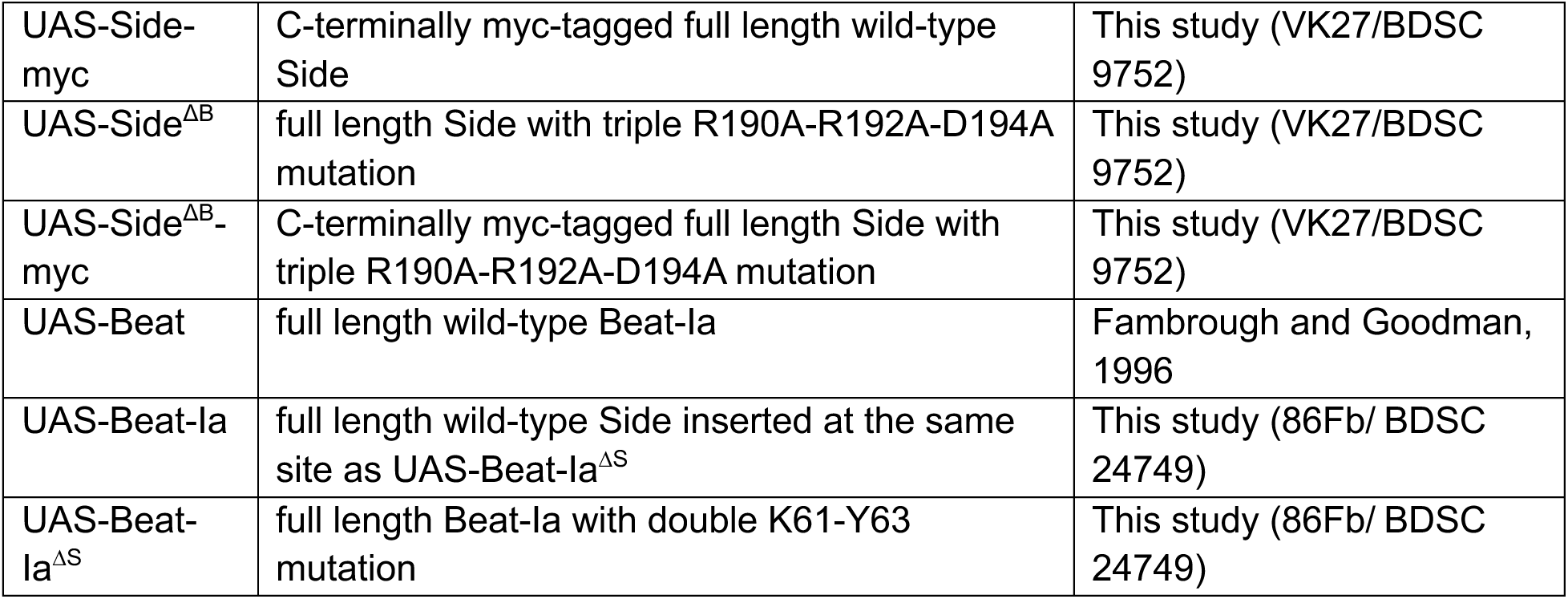

*Drosophila reagents* The following lines were used in this study:
| Genotype | Description | Source |
| --- | --- | --- |
| *side*<sup>ΔB</sup> | *side* allele with triple R190A-R192A-D194A mutation | This study |
| *side*<sup>I1563</sup> | null *side* allele, with shGFP background | Heymann et al., 2022 |
| *side*<sup>C137</sup> | null *side* allele | Heymann et al., 2022 |
| w1118 | white control | Bloomington Drosophila Stock Center (BDSC) |
| Mef2-GAL4 | Muscle GAL4 driver | Ranganayakulu et al., 1998 |
| UAS-Side | full length wild-type Side | This study (VK27/BDSC 9752) |
| UAS-Side-myc | C-terminally myc-tagged full length wild-type Side | This study (VK27/BDSC 9752) |
| UAS-Side<sup>ΔB</sup> | full length Side with triple R190A-R192A-D194A mutation | This study (VK27/BDSC 9752) |
| UAS-Side<sup>ΔB</sup>-myc | C-terminally myc-tagged full length Side with triple R190A-R192A-D194A mutation | This study (VK27/BDSC 9752) |
| UAS-Beat | full length wild-type Beat-Ia | Fambrough and Goodman, 1996 |
| UAS-Beat-Ia | full length wild-type Side inserted at the same site as UAS-Beat-Ia<sup>ΔS</sup> | This study (86Fb/ BDSC 24749) |
| UAS-Beat-Ia<sup>ΔS</sup> | full length Beat-Ia with double K61-Y63 mutation | This study (86Fb/ BDSC 24749) |

| Antibody | Concentration | Source |
| --- | --- | --- |
| Mouse anti-DLG | 1:100 | DSHB (4F3) |
| Rabbit anti-DLG | 1:40,000 | Koh et al., 1999 (PMID 10458610) |
| Mouse anti-Myc | 1:300 | Millipore (05-724) |
| Goat anti-Mouse 488 | 1:500 | Invitrogen (A11029) |
| Goat anti-Mouse 568 | 1:500 | Invitrogen (A11031) |
| Goat anti-Rabbit 488 | 1:500 | Invitrogen (A11008) |

### Drosophila dissections and immunohistochemistry

Third instar larval dissections and immunostaining were performed as previously described (Lobb-Rabe, Nawrocka, et al., 2024). Briefly, crawling third instar larvae were washed in deionized water and then dissected along the dorsal midline in phosphate-buffered saline (PBS, pH 7.4) on Sylgard dishes. After being fully stretched, dissected body walls were washed once with PBS and then fixed with 4% paraformaldehyde for 30 minutes. Larval fillets were washed in chilled PBST (PBS + 0.05% Triton X-100, pH 7.4) three times for 10 minutes on a nutator.

Samples were incubated in primary antibody at 4°C overnight. The antibodies used in this study and their working concentrations are listed in the table below. After three washes in PBST, samples were incubated in secondary antibody at room temperature for 2 hours. Samples were washed two times with PBST and once with PBS, then mounted in Vectashield (Vector Laboratories). Representative images were collected with on a Zeiss LSM800 confocal microscope with a 40× Plan-Neofluar 1.3NA objective and 0.5x zoom factor.

### Quantification and statistical analysis

Quantification of the innervation phenotype was conducted using a Zeiss AxioImager M2 with a Plan Neofluar 40x plan-neofluar 1.3NA objective. Connectivity was examined using DLG staining. Statistical analysis was performed using GraphPad (Prism) and SciPy (version 1.5.2, Anaconda). Each genotype was collected from a minimum of two separate crosses, and connectivity on intact muscles from abdominal body wall segments 2–5 was recorded. p-value and significance were determined through Chi Squared Test with Fisher’s exact test; see Table S3 for a list of results of significance tests for genotypes tested in the manuscript.

## SUPPLEMENTAL FIGURE LEGENDS

**Figure S1.**
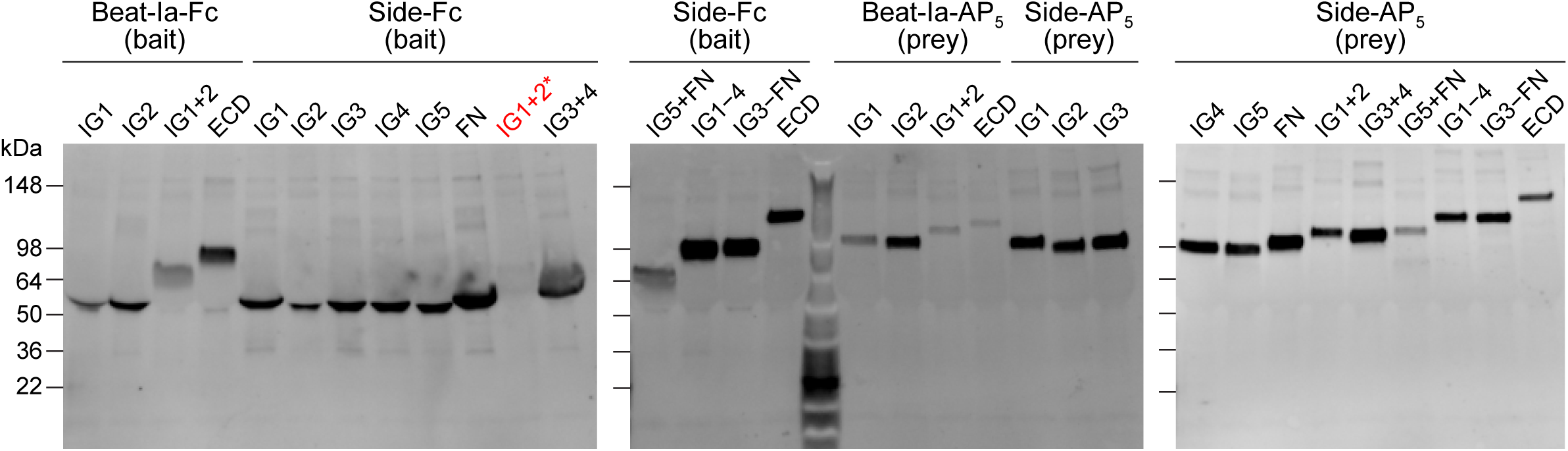
Protein expression levels for ECIA constructs in Figure 1. Anti-His western blots of Beat-Ia and Side domains in bait and prey form. The Side-Fc IG1+2 construct showed no expression (labeled red).

**Figure S2.**
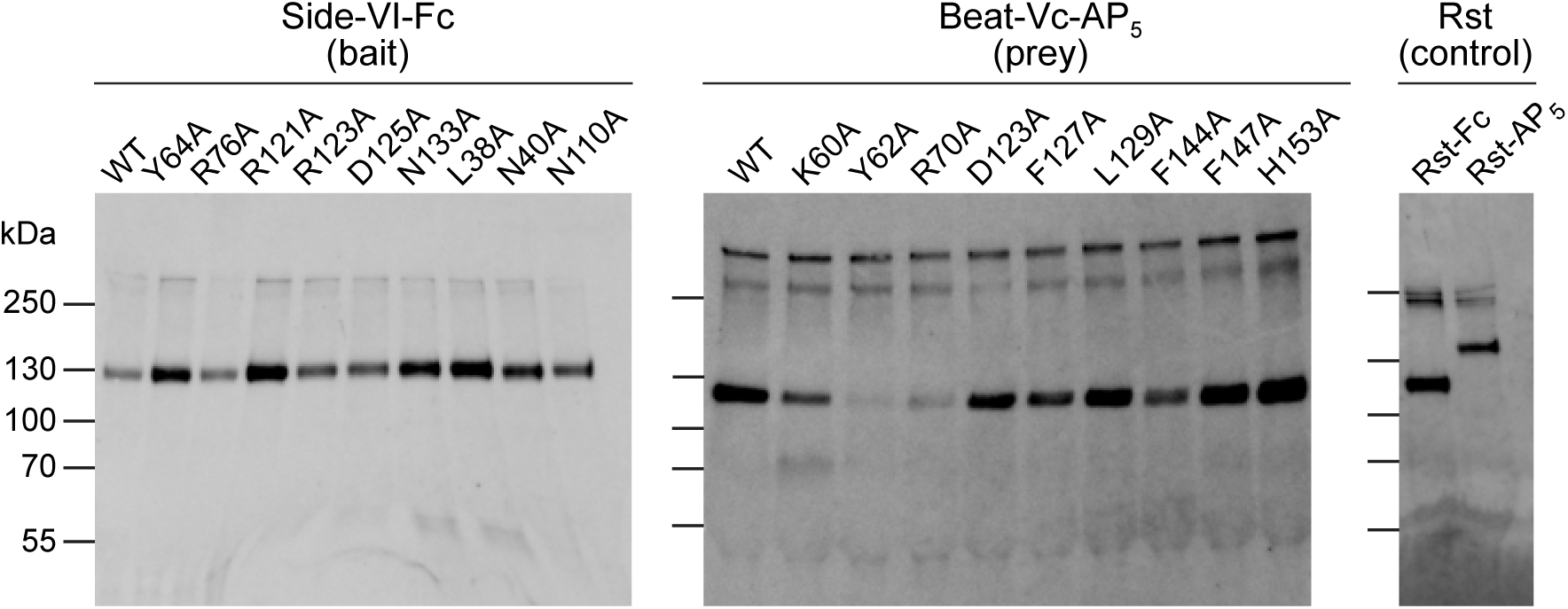
Protein expression levels for ECIA constructs in Figure 2. Anti-His western blots of Beat-Vc and Side-VI mutants in prey and bait form, respectively. The five-IG domain *Drosophila* Rst protein is used as a negative control against Beats and Sides, while the homodimeric Rst-Rst interaction serves as a positive control for the assay.

**Figure S3.**
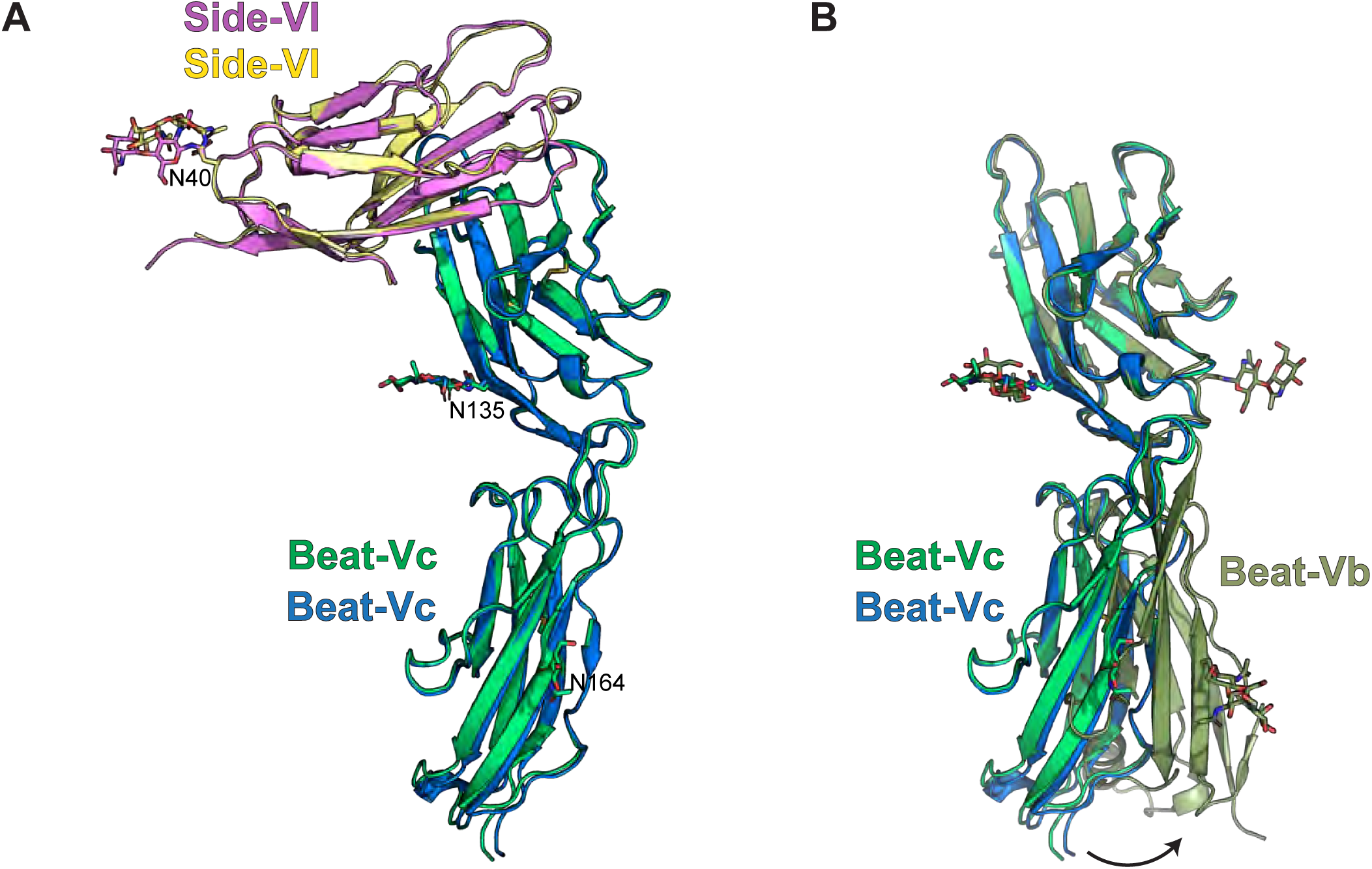
Flexibility in the Beat IG-IG domain boundary. **A.** The two copies of the Beat-Vc IG1+2–Side-VI IG1 complex in the asymmetric unit show no significant differences. **B.** Comparison of our Beat-Vb structure with Beat-Vc from the complex crystals show variability in the orientation of Beat IG1 and IG2 domains with respect to each other.

**Figure S4.**
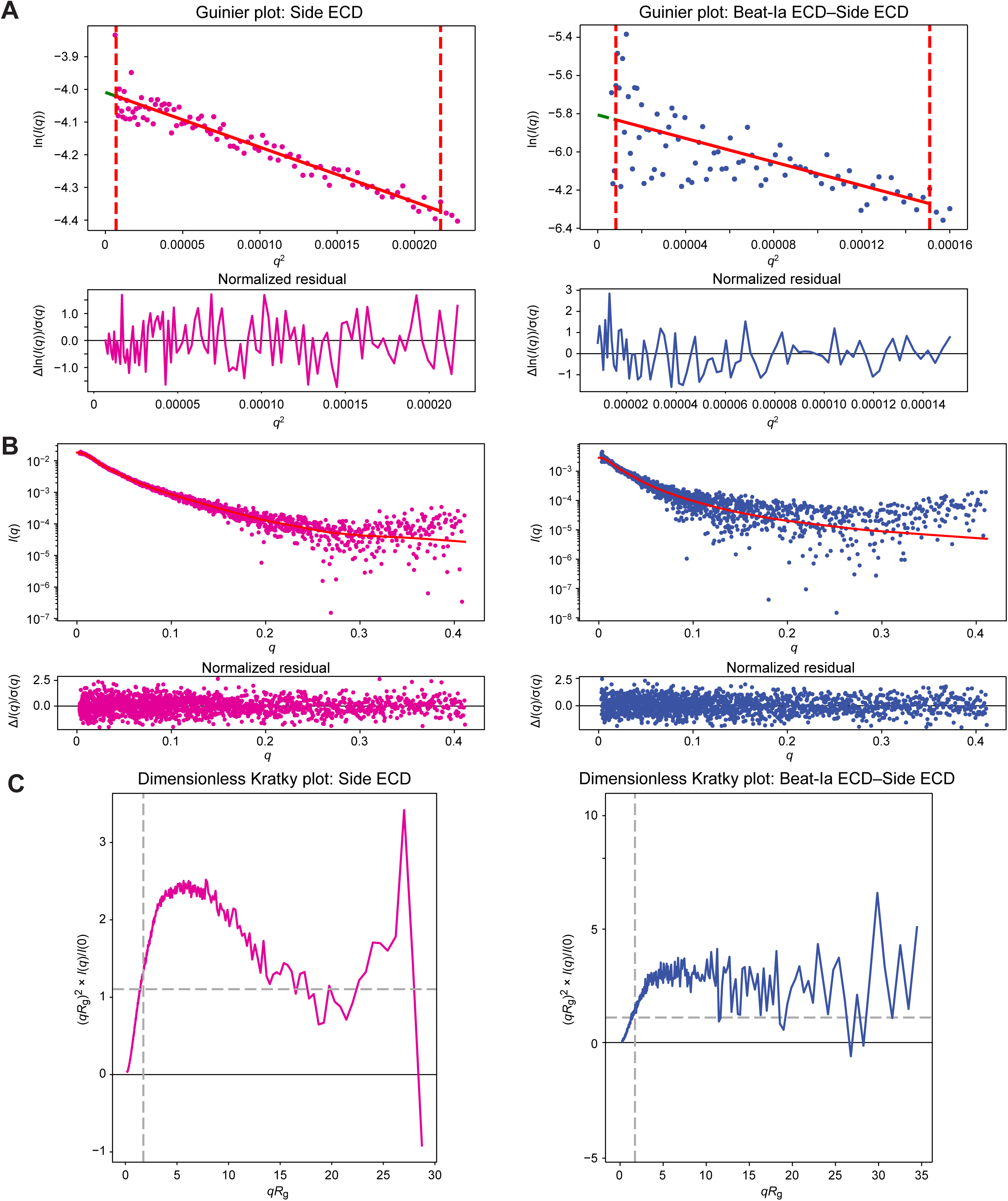
Small-angle x-ray scattering analysis **A.** Guinier plots for Side ECD (left) and Beat-Ia ECD–Side ECD complex (right). **B.** Indirect Fourier Transform fit to scattering profiles, as performed by GNOM in RAW. P(*r*) plots are shown in Figures 5C and 5E. **C.** Dimensionless Kratky plots. The dashed lines label (√3,3/e).

**Figure S5.**
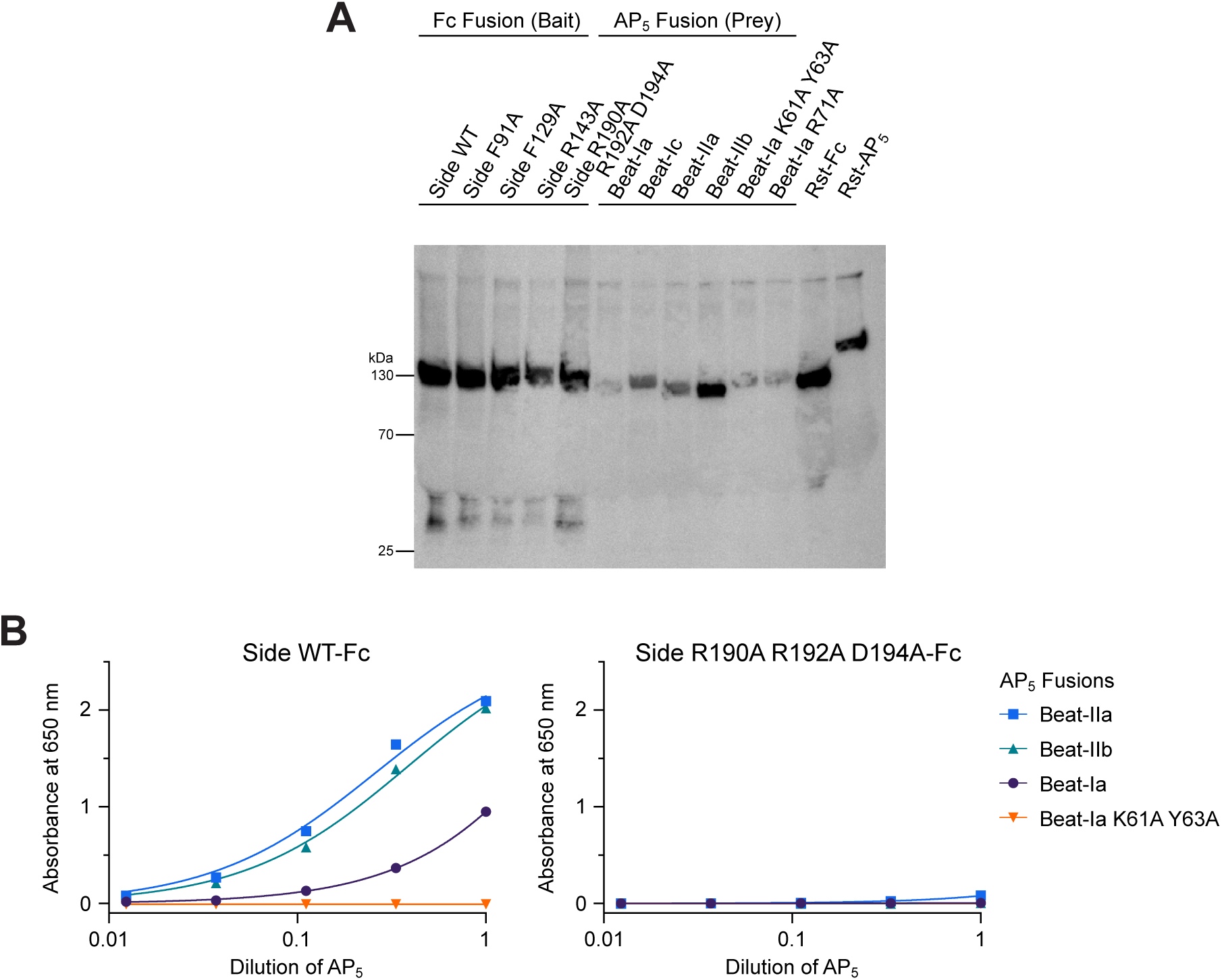
Beat-Ia K61A-Y63A and Side R190A-R192A-D194A do not interact with wild-type Side and Beat-Ia, respectively. **A.** Anti-His western blots of Beat and Side mutants used for ECIA experiments in Figure 5. **B.** Plot of ECIA absorbance values in Figure 5F, fit to a binding isotherm.

**Figure S6.**
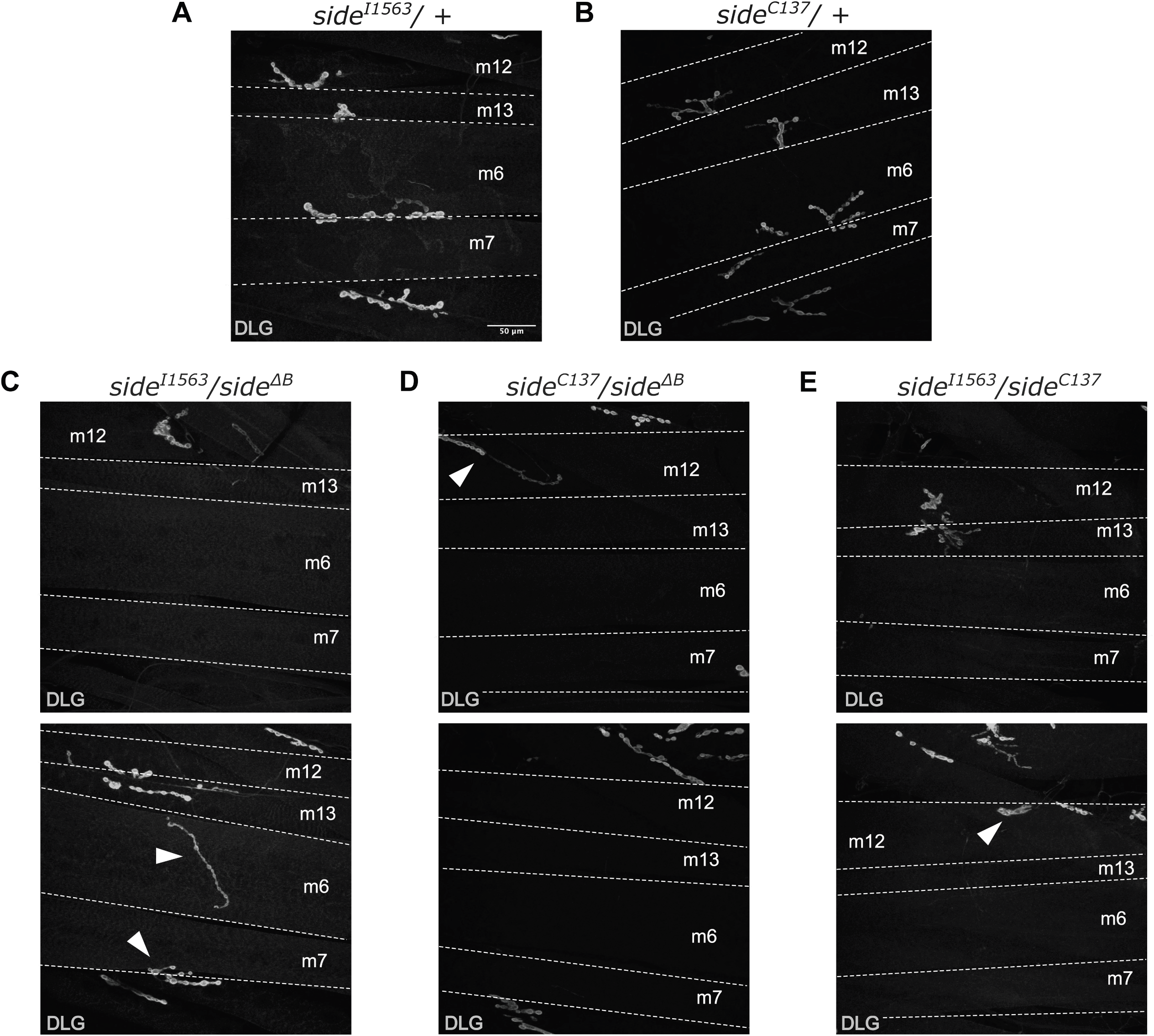
Larvae with different *side* alleles display a variety of connectivity patterns. **A.** *side^I1563^*/+ animals display normal innervation of ventral muscles. **B.** *side^C137^*/+ animals display normal innervation of ventral muscles. **C.** *side^I1563^*/*side^ΔB^* animals display loss of innervation on a subset of muscles (top) and ectopic innervations (bottom; arrowheads). **D.** *side^C137^*/*side^ΔB^* animals display loss and ectopic innervation on a subset of muscles (top; arrowhead) and loss of innervations (bottom). **E.** *side^I1563^*/*side^ΔB^* animals display loss of innervation (top) and ectopic innervation (bottom; arrowhead).

**Figure S7.**
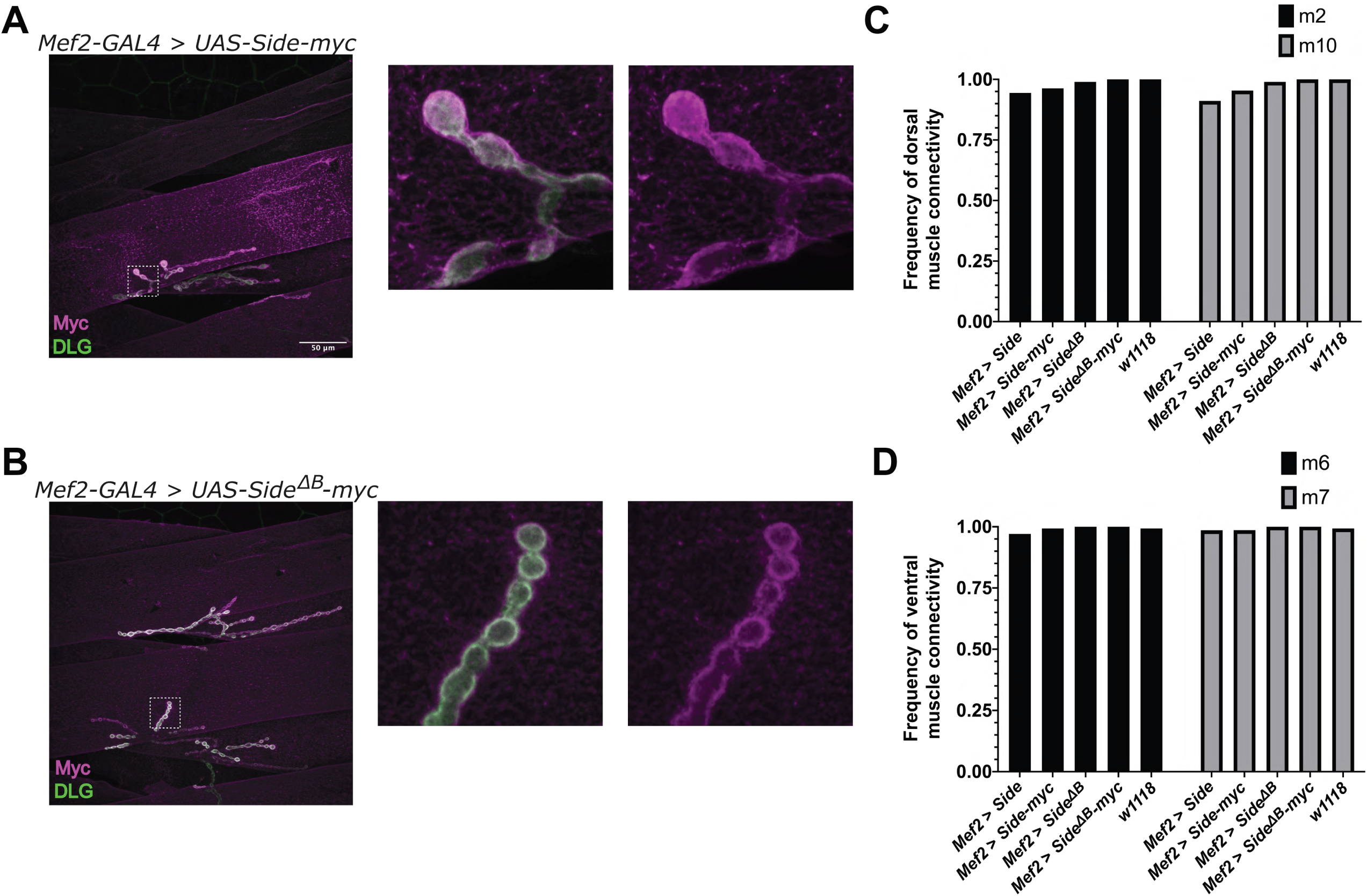
Overexpressed Side transgenes localize to the postsynaptic membrane and some muscles retain innervation. **A,B.** Representative images of dorsal muscles labeled with anti-DLG (green) and anti-Myc (magenta). Dashed boxes refer to the zoomed in region (right) to more clearly show localization of Side and Side^ΔB^ to the postsynaptic membrane. **C.** Quantification of connectivity frequency on muscle 2 and 10. (n = 14-24) **D.** Quantification of connectivity frequency on muscle 6 and 7. (Sample size: *Mef2 > Side*, n = 22; *Mef2 > Side-myc*, n = 24; *Mef2 >Side^ΔB^*, n = 22; *Mef2 > Side^ΔB^-myc*, n = 14; *w^1118^*, n = 18)

## SUPPLEMENTAL TABLES

**Table S1.** Data and refinement statistics for x-ray crystallography.

|  | Beat-Vb<br>IG1+2 | Side-IV IG1<br>(Se-Met labeled) | Side-VI IG1<br>(refolded) | Side-VII IG1 | Beat-Vc IG1+2<br>+ Side-VI IG1 |
| --- | --- | --- | --- | --- | --- |
| **Data Collection** |  |  |  |  |  |
| Beamline | APS 23-ID-B | APS 24-ID-C | SSRL 12-2 | APS 23-ID-B | APS 23-ID-B |
| Space Group | *P*6<sub>2</sub>22 | *C*2 | *C*222<sub>1</sub> | *P*4<sub>3</sub>2<sub>1</sub>2 | *P*6<sub>1</sub> |
| *Cell Dimensions* |  |  |  |  |  |
| *a, b, c* (Å) | 183.68, 183.68, 68.41 | 94.54, 29.77, 41.97 | 72.77, 181.42, 300.12 | 37.38, 37.38, 189.66 | 87.48, 87.48, 387.38 |
| $\alpha, \beta, \gamma$ (°) | 90, 90, 120 | 90, 91.43, 90 | 90, 90, 90 | 90, 90, 90 | 90, 90, 120 |
| Resolution (Å) | 50-3.40 (3.49-3.40)* | 50-2.40 (2.54-2.40) | 50-2.51 (2.57-2.51) | 50-2.51 (2.71-2.51) | 50-3.93 (4.20-3.93) |
| *R*<sub>sym</sub> (%) | 15.8 (337.8) | 8.3 (25.4)*** | 10.4 (112.1) | 9.9 (68.0) | 23.8 (50.0) |
| *R*<sub>meas</sub> (%) | 16.0 (330.6) | 11.5 (35.0) | 11.3 (128.5) | 10.8 (77.9) | 25.6 (53.8) |
| $\langle I/\sigma(I) \rangle$ | 18.82 (1.71) | 7.87 (3.40) | 15.66 (1.22) | 12.16 (2.30) | 5.99 (3.82) |
| *CC*<sub>1/2</sub> | 0.994 (0.402) | 0.987 (0.857) | 0.998 (0.547) | 0.998 (0.711) | 0.991 (0.975) |
| Completeness (%) (spherical) | 99.4 (95.5) | 86.6 (81.7) | 97.8 (73.8) | 70.6 (36.3) | 64.9 (26.0) |
| (ellipsoidal) | N/A | N/A | N/A | 92.5 (83.2)<sup>†</sup> | 91.5 (99.1)<sup>††</sup> |
| Redundancy | 40.9 (34.7) | 1.6 (1.5) | 6.3 (3.0) | 6.3 (4.0) | 7.6 (7.3) |
| **Refinement** |  |  |  |  |  |
| Resolution (Å) | 50-3.40 (3.89-3.40)* | 50-2.40 (2.74-2.40) | 50-2.51 (2.57-2.51) | 50-2.51 | 50-3.93 (4.50-3.93) |
| Reflections | 9,745 | 4,533 | 66,830 | 3,659 | 9,644 |
| *R*<sub>cryst</sub> (%) | 29.63 (36.54) | 18.83 (20.81) | 21.24 (41.08) | 24.64 | 23.38 (26.80) |
| *R*<sub>free</sub> (%)** | 31.50 (43.61) | 22.04 (27.00) | 23.70 (44.72) | 28.02 | 28.37 (31.35) |
| *Number of atoms* |  |  |  |  |  |
| Protein | 1977 | 925 | 9282 | 834 | 5516 |
| Ligand/Glycans | 95 | 6 | 16 | 0 | 112 |
| Water | 0 | 39 | 121 | 0 | 0 |
| *Average B-factors* (Å<sup>2</sup>) |  |  |  |  |  |
| All | 205.1 | 30.7 | 63.5 | 64.8 | 81.8 |
| Protein | 204.5 | 30.8 | 63.6 | 64.8 | 81.0 |
| Ligand/Glycans | 217.9 | 36.1 | 64.8 | N/A | 118.5 |
| Solvent | N/A | 27.4 | 55.8 | N/A | N/A |
| *R.m.s. deviations from ideality* |  |  |  |  |  |
| Bond Lengths (Å) | 0.004 | 0.005 | 0.003 | 0.007 | 0.004 |
| Bond Angles (°) | 0.669 | 0.727 | 0.531 | 0.894 | 0.771 |
| *Ramachandran plot* |  |  |  |  |  |
| Favored (%) | 93.83 | 96.46 | 97.20 | 92.63 | 93.15 |
| Outliers (%) | 0.00 | 0.00 | 0.00 | 0.00 | 0.00 |
| Rotamer Outliers (%) | 2.23 | 1.98 | 1.43 | 4.26 | 0.33 |
| All-atom Clashscore<sup>‡</sup> | 13.96 | 3.80 | 5.74 | 7.27 | 9.78 |

\* The values in parentheses are for reflections in the highest resolution bin.
\*\* 5% of reflections (488) for Beat-Vb, 10% of reflections (453) for Side-IV, 3% of reflections (1,946) for Side-VI, 5% for Side-VII (184), and 5% of reflections (455) for Beat-Vc–Side-VI datasets were not used during refinement for cross validation.
<sup>†</sup> Diffraction limits (Å) and corresponding principal axes of the ellipsoid fitted to the diffraction cut-off surface as direction cosines in the orthogonal basis (standard PDB convention), and in terms of reciprocal unit-cell vectors, are:
Diffraction limit #1: 4.972 Å 1.0000 0.0000 0.0000 (0.894 **a**\* – 0.447 **b**\*) Diffraction limit #2: 4.972 Å 0.0000 1.0000 0.0000 **b**\* Diffraction limit #3: 3.147 Å 0.0000 0.0000 1.0000 **c**\*
<sup>††</sup> Diffraction limits (Å) and corresponding principal axes of the ellipsoid fitted to the diffraction cut-off surface as direction cosines in the orthogonal basis (standard PDB convention), and in terms of reciprocal unit-cell vectors, are:
Diffraction limit #1: 2.273 Å 1.0000 0.0000 0.0000 **a**\* Diffraction limit #2: 2.273 Å 0.0000 1.0000 0.0000 **b**\* Diffraction limit #3: 3.591 Å 0.0000 0.0000 1.0000 **c**\*
\*\*\* Scaled with Friedel pairs separated.
<sup>‡</sup> Clashscores were calculated by *phenix.refine* (Phenix version 1.21).
N/A: Not applicable.

**Table S2.** Data statistics and analysis for small angle x-ray scattering experiments.

|  | Side ECD | Side ECD–Beat-Ia ECD<br>(1:1.3 molar mix) |
| --- | --- | --- |
| *Data Collection and Sample Details* |  |  |
| Experiment setup | SEC-MALS-SAXS | SEC-MALS-SAXS |
| Instrument | BioCAT facility at APS beamline 18-ID with Eiger2 XE 9M detector with buffer co-flow during SAXS |  |
| Wavelength (Å) | 1.033 | 1.033 |
| Camera length (m) | 3.725 | 3.725 |
| $q$-measurement range (Å<sup>-1</sup>) | 0.0025 to 0.41 | 0.0025 to 0.41 |
| Exposure time (s) | 0.15 | 0.15 |
| Exposure period (s) | 1.0 | 1.0 |
| Flow rate (ml/min) | 0.6 | 0.45 |
| Chromatography column | Superdex 200 10/300 Increase |  |
| Buffer | 10 mM HEPES pH 7.4,<br>150 mM NaCl | 10 mM HEPES pH 7.2,<br>500 mM NaCl, 10%<br>Glycerol |
| Temperature | 22°C | 22°C |
| Software | BioXTAS RAW version 2.3.0, ATSAS 4.0.1 |  |
| Loading concentration (mg/ml) | 3.0 | 1.5 |
| Loading volume (μl) | 230 | 230 |
| *Structural Parameters* |  |  |
| Guinier Analysis* |  |  |
| $I(0)$ (cm<sup>-1</sup>) | 0.018 ± 0.00011 | 0.0030 ± 0.000098 |
| $R_g$ (Å) | 70.90 ± 1.04 | 96.14 ± 5.66 |
| $q$ range (Å<sup>-1</sup>) | 0.00263 to 0.01473 | 0.00287 to 0.01228 |
| $q_{\min}R_g$ to $q_{\max}R_g$ | 0.1863 to 1.0443 | 0.2762 to 1.1810 |
| Coefficient of Correlation, $R^2$ | 0.931 | 0.473 |
| $P(r)$ Analysis (from GNOM)** | | |
| $I(0)$ (cm<sup>-1</sup>) | 0.018 ± 0.00011 | 0.0029 ± 0.00007 |
| $R_g$ (Å) | 73.75 ± 0.87 | 91.06 ± 3.35 |
| $D_{\max}$ (Å) | 281 | 333 |
| $q$ range (Å<sup>-1</sup>) | 0.0026 to 0.4114 | 0.0029 to 0.411 |
| $\chi^2$ | 0.5547 | 0.5275 |
| Molecular Weight Estimates |  |  |
| Vc method*** | 87.4 kDa | 108.0 kDa |
| Vp method† | 195.1 kDa | 305.7 MDa |
| Shape & Size method‡ | extended | extended |

\* Performed in BioXTAS RAW (42).
\*\* By Svergun (43), as used within RAW.
\*\*\* By Rambo and Tainer (45). $q_{\max}$ was set to 0.3.
† By Piiadov, et al. (46). This method does not work well with non-globular or flexible samples and is likely not appropriate in this case.
‡ By Franke, et al. (47). This method may not work well with non-protein samples.

**Table S3.** Results of statistical testing performed in the manuscript.

| **Genotype 1** | **Genotype 2** | **Muscle** | **Fisher p-value** | **Statistically Significant**<br>**(P < 0.05)** |
| --- | --- | --- | --- | --- |
| *side*<sup>ΔB</sup> | *side*<sup>ΔB</sup> / *side*<sup>I1563</sup> | m6 | 0.012549169 | Yes |
| *side*<sup>ΔB</sup> | *side*<sup>ΔB</sup> / *side*<sup>C137</sup> | m6 | 6.88282E-05 | Yes |
| *side*<sup>ΔB</sup> | *side*<sup>I1563</sup> / *side*<sup>C137</sup> | m6 | 2.62736E-11 | Yes |
| *side*<sup>ΔB</sup> | *side*<sup>ΔB</sup> / + | m6 | 2.54647E-09 | Yes |
| *side*<sup>ΔB</sup> | *side*<sup>I1563</sup> / + | m6 | 1.63174E-10 | Yes |
| *side*<sup>ΔB</sup> | *side*<sup>C137</sup> / + | m6 | 6.13704E-07 | Yes |
| *side*<sup>ΔB</sup> / *side*<sup>I1563</sup> | *side*<sup>ΔB</sup> / *side*<sup>C137</sup> | m6 | 0.157469569 | No |
| *side*<sup>ΔB</sup> / *side*<sup>I1563</sup> | *side*<sup>I1563</sup> / *side*<sup>C137</sup> | m6 | 3.37253E-05 | Yes |
| *side*<sup>ΔB</sup> / *side*<sup>I1563</sup> | *side*<sup>ΔB</sup> / + | m6 | 2.22617E-16 | Yes |
| *side*<sup>ΔB</sup> / *side*<sup>I1563</sup> | *side*<sup>I1563</sup> / + | m6 | 8.02202E-18 | Yes |
| *side*<sup>ΔB</sup> / *side*<sup>I1563</sup> | *side*<sup>C137</sup> / + | m6 | 3.09164E-13 | Yes |
| *side*<sup>ΔB</sup> / *side*<sup>C137</sup> | *side*<sup>ΔB</sup> / *side*<sup>C137</sup> | m6 | 0.004829352 | Yes |
| *side*<sup>ΔB</sup> / *side*<sup>C137</sup> | *side*<sup>ΔB</sup> / + | m6 | 7.45476E-22 | Yes |
| *side*<sup>ΔB</sup> / *side*<sup>C137</sup> | *side*<sup>I1563</sup> / + | m6 | 4.67669E-23 | Yes |
| *side*<sup>ΔB</sup> / *side*<sup>C137</sup> | *side*<sup>C137</sup> / + | m6 | 9.04515E-18 | Yes |
| *side*<sup>I1563</sup> / *side*<sup>C137</sup> | *side*<sup>ΔB</sup> / + | m6 | 2.92761E-33 | Yes |
| *side*<sup>I1563</sup> / *side*<sup>C137</sup> | *side*<sup>I1563</sup> / + | m6 | 9.40443E-35 | Yes |
| *side*<sup>I1563</sup> / *side*<sup>C137</sup> | *side*<sup>C137</sup> / + | m6 | 2.21657E-28 | Yes |
| *side*<sup>ΔB</sup> / + | *side*<sup>I1563</sup> / + | m6 | 1 | No |
| *side*<sup>ΔB</sup> / + | *side*<sup>C137</sup> / + | m6 | 0.350383177 | No |
| *side*<sup>I1563</sup> / + | *side*<sup>C137</sup> / + | m6 | 0.109889939 | No |
| *side*<sup>ΔB</sup> | *side*<sup>ΔB</sup> / *side*<sup>I1563</sup> | m7 | 0.001278148 | Yes |
| *side*<sup>ΔB</sup> | *side*<sup>ΔB</sup> / *side*<sup>C137</sup> | m7 | 2.6299E-06 | Yes |
| *side*<sup>ΔB</sup> | *side*<sup>I1563</sup> / *side*<sup>C137</sup> | m7 | 3.17197E-11 | Yes |
| *side*<sup>ΔB</sup> | *side*<sup>ΔB</sup> / + | m7 | 2.33451E-09 | Yes |
| *side*<sup>ΔB</sup> | *side*<sup>I1563</sup> / + | m7 | 1.21365E-11 | Yes |
| *side*<sup>ΔB</sup> | *side*<sup>C137</sup> / + | m7 | 6.8509E-08 | Yes |
| *side*<sup>ΔB</sup> / *side*<sup>I1563</sup> | *side*<sup>ΔB</sup> / *side*<sup>C137</sup> | m7 | 0.158448387 | No |
| *side*<sup>ΔB</sup> / *side*<sup>I1563</sup> | *side*<sup>I1563</sup> / *side*<sup>C137</sup> | m7 | 0.000673163 | Yes |
| *side*<sup>ΔB</sup> / *side*<sup>I1563</sup> | *side*<sup>ΔB</sup> / + | m7 | 3.13978E-19 | Yes |
| *side*<sup>ΔB</sup> / *side*<sup>I1563</sup> | *side*<sup>I1563</sup> / + | m7 | 6.99334E-22 | Yes |
| *side*<sup>ΔB</sup> / *side*<sup>I1563</sup> | *side*<sup>C137</sup> / + | m7 | 7.4912E-17 | Yes |
| *side*<sup>ΔB</sup> / *side*<sup>C137</sup> | *side*<sup>I1563</sup> / *side*<sup>C137</sup> | m7 | 0.049419421 | Yes |
| *side*<sup>ΔB</sup> / *side*<sup>C137</sup> | *side*<sup>ΔB</sup> / + | m7 | 2.69766E-25 | Yes |
| *side*<sup>ΔB</sup> / *side*<sup>C137</sup> | *side*<sup>I1563</sup> / + | m7 | 5.63119E-28 | Yes |
| *side*<sup>ΔB</sup> / *side*<sup>C137</sup> | *side*<sup>C137</sup> / + | m7 | 2.31668E-22 | Yes |
| *side*<sup>I1563</sup> / *side*<sup>C137</sup> | *side*<sup>ΔB</sup> / + | m7 | 1.60739E-33 | Yes |
| *side*<sup>I1563</sup> / *side*<sup>C137</sup> | *side*<sup>I1563</sup> / + | m7 | 1.36372E-36 | Yes |
| *side*<sup>I1563</sup> / *side*<sup>C137</sup> | *side*<sup>C137</sup> / + | m7 | 4.00188E-30 | Yes |
| *side*<sup>ΔB</sup> / + | *side*<sup>I1563</sup> / + | m7 | 0.498024096 | No |
| *side*<sup>ΔB</sup> / + | *side*<sup>C137</sup> / + | m7 | 0.671637581 | No |
| *side*<sup>I1563</sup> / + | *side*<sup>C137</sup> / + | m7 | 0.109889939 | No |
| *side*<sup>ΔB</sup> | *side*<sup>ΔB</sup> / *side*<sup>I1563</sup> | m12 | 0.019884 | Yes |
| $side^{\Delta B}$ | $side^{\Delta B} / side^{C137}$ | m12 | 0.864316 | No |
| $side^{\Delta B}$ | $side^{I1563} / side^{C137}$ | m12 | 0.323816 | No |
| $side^{\Delta B}$ | $side^{\Delta B} / +$ | m12 | 0.00000694 | Yes |
| $side^{\Delta B}$ | $side^{I1563} / +$ | m12 | 0.0000506 | Yes |
| $side^{\Delta B}$ | $side^{C137} / +$ | m12 | 0.000125 | Yes |
| $side^{\Delta B} / side^{I1563}$ | $side^{\Delta B} / side^{C137}$ | m12 | 0.032197 | Yes |
| $side^{\Delta B} / side^{I1563}$ | $side^{I1563} / side^{C137}$ | m12 | 0.23347 | No |
| $side^{\Delta B} / side^{I1563}$ | $side^{\Delta B} / +$ | m12 | 3.27E-11 | Yes |
| $side^{\Delta B} / side^{I1563}$ | $side^{I1563} / +$ | m12 | 4.78E-10 | Yes |
| $side^{\Delta B} / side^{I1563}$ | $side^{C137} / +$ | m12 | 1.9E-09 | Yes |
| $side^{\Delta B} / side^{C137}$ | $side^{I1563} / side^{C137}$ | m12 | 0.420378 | No |
| $side^{\Delta B} / side^{C137}$ | $side^{\Delta B} / +$ | m12 | 0.00000228 | Yes |
| $side^{\Delta B} / side^{C137}$ | $side^{I1563} / +$ | m12 | 0.0000309 | Yes |
| $side^{\Delta B} / side^{C137}$ | $side^{C137} / +$ | m12 | 0.0000412 | Yes |
| $side^{I1563} / side^{C137}$ | $side^{\Delta B} / +$ | m12 | 4.05E-08 | Yes |
| $side^{I1563} / side^{C137}$ | $side^{I1563} / +$ | m12 | 0.00000041 | Yes |
| $side^{I1563} / side^{C137}$ | $side^{C137} / +$ | m12 | 0.00000115 | Yes |
| $side^{\Delta B} / +$ | $side^{I1563} / +$ | m12 | 0.620482 | No |
| $side^{\Delta B} / +$ | $side^{C137} / +$ | m12 | 0.606927 | No |
| $side^{I1563} / +$ | $side^{C137} / +$ | m12 | 1 | No |
| $side^{\Delta B}$ | $side^{\Delta B} / side^{I1563}$ | m13 | 0.000439184 | Yes |
| $side^{\Delta B}$ | $side^{\Delta B} / side^{C137}$ | m13 | 0.003119584 | Yes |
| $side^{\Delta B}$ | $side^{I1563} / side^{C137}$ | m13 | 8.77828E-10 | Yes |
| $side^{\Delta B}$ | $side^{\Delta B} / +$ | m13 | 2.25507E-05 | Yes |
| $side^{\Delta B}$ | $side^{I1563} / +$ | m13 | 0.000122456 | Yes |
| $side^{\Delta B}$ | $side^{C137} / +$ | m13 | 0.002584192 | Yes |
| $side^{\Delta B} / side^{I1563}$ | $side^{\Delta B} / side^{C137}$ | m13 | 0.594519932 | No |
| $side^{\Delta B} / side^{I1563}$ | $side^{I1563} / side^{C137}$ | m13 | 0.006933506 | Yes |
| $side^{\Delta B} / side^{I1563}$ | $side^{\Delta B} / +$ | m13 | 6.90502E-14 | Yes |
| $side^{\Delta B} / side^{I1563}$ | $side^{I1563} / +$ | m13 | 9.36461E-13 | Yes |
| $side^{\Delta B} / side^{I1563}$ | $side^{C137} / +$ | m13 | 1.82949E-10 | Yes |
| $side^{\Delta B} / side^{C137}$ | $side^{I1563} / side^{C137}$ | m13 | 0.001332665 | Yes |
| $side^{\Delta B} / side^{C137}$ | $side^{\Delta B} / +$ | m13 | 1.09541E-12 | Yes |
| $side^{\Delta B} / side^{C137}$ | $side^{I1563} / +$ | m13 | 2.35779E-11 | Yes |
| $side^{\Delta B} / side^{C137}$ | $side^{C137} / +$ | m13 | 3.07286E-09 | Yes |
| $side^{I1563} / side^{C137}$ | $side^{\Delta B} / +$ | m13 | 1.12761E-23 | Yes |
| $side^{I1563} / side^{C137}$ | $side^{I1563} / +$ | m13 | 8.09109E-22 | Yes |
| $side^{I1563} / side^{C137}$ | $side^{C137} / +$ | m13 | 6.10524E-19 | Yes |
| $side^{\Delta B} / +$ | $side^{I1563} / +$ | m13 | 0.68120022 | No |
| $side^{\Delta B} / +$ | $side^{C137} / +$ | m13 | 0.263137798 | No |
| $side^{I1563} / +$ | $side^{C137} / +$ | m13 | 0.487818871 | No |

| Genotype 1 | Genotype 2 | Muscle | Fisher p-value | Statistically Significant (P < 0.05) |
| --- | --- | --- | --- | --- |
| *Mef2* > *Side* | *Mef2* > *Side-myc* | m1 | 0.199361737 | No |
| *Mef2* > *Side* | *Mef2* > *Side*<sup>ΔB</sup> | m1 | 8.36113E-17 | Yes |
| *Mef2* > *Side* | *Mef2* > *Side*<sup>ΔB</sup> -myc | m1 | 1.2144E-14 | Yes |
| *Mef2* > *Side* | *w*<sup>1118</sup> | m1 | 6.47994E-18 | Yes |
| *Mef2* > *Side-myc* | *Mef2* > *Side*<sup>ΔB</sup> | m1 | 1.42805E-13 | Yes |
| *Mef2* > *Side-myc* | *Mef2* > *Side*<sup>ΔB</sup> -myc | m1 | 1.34551E-11 | Yes |
| *Mef2* > *Side-myc* | *w*<sup>1118</sup> | m1 | 9.56977E-15 | Yes |
| *Mef2* > *Side*<sup>ΔB</sup> | *Mef2* > *Side*<sup>ΔB</sup> -myc | m1 | 1 | No |
| *Mef2* > *Side*<sup>ΔB</sup> | *w*<sup>1118</sup> | m1 | 0.499764982 | No |
| *Mef2* > *Side*<sup>ΔB</sup> -myc | *w*<sup>1118</sup> | m1 | 0.236740804 | No |
| *Mef2* > *Side* | *Mef2* > *Side-myc* | m9 | 0.198327114 | No |
| *Mef2* > *Side* | *Mef2* > *Side*<sup>ΔB</sup> | m9 | 3.06843E-21 | Yes |
| *Mef2* > *Side* | *Mef2* > *Side*<sup>ΔB</sup> -myc | m9 | 1.1034E-18 | Yes |
| *Mef2* > *Side* | *w*<sup>1118</sup> | m9 | 6.79631E-21 | Yes |
| *Mef2* > *Side-myc* | *Mef2* > *Side*<sup>ΔB</sup> | m9 | 1.08032E-17 | Yes |
| *Mef2* > *Side-myc* | *Mef2* > *Side*<sup>ΔB</sup> -myc | m9 | 1.99353E-15 | Yes |
| *Mef2* > *Side-myc* | *w*<sup>1118</sup> | m9 | 2.46387E-17 | Yes |
| *Mef2* > *Side*<sup>ΔB</sup> | *Mef2* > *Side*<sup>ΔB</sup> -myc | m9 | 1 | No |
| *Mef2* > *Side*<sup>ΔB</sup> | *w*<sup>1118</sup> | m9 | 1 | No |
| *Mef2* > *Side*<sup>ΔB</sup> -myc | *w*<sup>1118</sup> | m9 | 0.488095238 | No |
| *Mef2* > *Side-myc, Beat-Ia* | *Mef2* > *Side-myc, Beat-Ia*<sup>ΔS</sup> | m1 | 0.166308069 | No |
| *Mef2* > *Side-myc, Beat-Ia* | *Mef2* > *Side-myc* | m1 | 0.061756633 | No |
| *Mef2* > *Side-myc, Beat-Ia*<sup>ΔS</sup> | *Mef2* > *Side-myc* | m1 | 0.75402109 | No |
| *Mef2* > *Side-myc, Beat-Ia* | *Mef2* > *Side-myc, Beat-Ia*<sup>ΔS</sup> | m9 | 0.003200174 | Yes |
| *Mef2* > *Side-myc, Beat-Ia* | *Mef2* > *Side-myc* | m9 | 0.045197039 | Yes |
| *Mef2* > *Side-myc, Beat-Ia*<sup>ΔS</sup> | *Mef2* > *Side-myc* | m9 | 0.289800886 | No |

